# Transcription factor NF-κB in a basal metazoan, the sponge, has conserved and unique sequences, activities, and regulation

**DOI:** 10.1101/691097

**Authors:** Leah M. Williams, Melissa M. Inge, Katelyn M. Mansfield, Anna Rasmussen, Jamie Afghani, Mikhail Agrba, Colleen Albert, Cecilia Andersson, Milad Babaei, Mohammad Babaei, Abigail Bagdasaryants, Arianna Bonilla, Amanda Browne, Sheldon Carpenter, Tiffany Chen, Blake Christie, Andrew Cyr, Katie Dam, Nicholas Dulock, Galbadrakh Erdene, Lindsie Esau, Stephanie Esonwune, Anvita Hanchate, Xinli Huang, Timothy Jennings, Aarti Kasabwala, Leanne Kehoe, Ryan Kobayashi, Migi Lee, Andre LeVan, Yuekun Liu, Emily Murphy, Avanti Nambiar, Meagan Olive, Devansh Patel, Flaminio Pavesi, Christopher A. Petty, Yelena Samofalova, Selma Sanchez, Camilla Stejskal, Yinian Tang, Alia Yapo, John P. Cleary, Sarah A. Yunes, Trevor Siggers, Thomas D. Gilmore

## Abstract

Biological and biochemical functions of immunity transcription factor NF-κB in basal metazoans are largely unknown. Herein, we characterize transcription factor NF-κB from the demosponge *Amphimedon queenslandica* (Aq), in the phylum Porifera. Structurally and phylogenetically, the Aq-NF-κB protein is most similar to NF-κB p100 and p105 among vertebrate proteins, with an N-terminal DNA-binding/dimerization domain, a C-terminal Ankyrin (ANK) repeat domain, and a DNA binding-site profile more similar to human NF-κB proteins than Rel proteins. Aq-NF-κB also resembles the mammalian NF-κB protein p100 in that C-terminal truncation results in translocation of Aq-NF-κB to the nucleus and increases its transcriptional activation activity. Overexpression of a human or sea anemone IκB kinase (IKK) can induce C-terminal processing of Aq-NF-κB *in vivo*, and this processing requires C-terminal serine residues in Aq-NF-κB. Unlike human NF-κB p100, however, the C-terminal sequences of Aq-NF-κB do not effectively inhibit its DNA-binding activity when expressed in human cells. Tissue of another demosponge, a black encrusting sponge, contains NF-κB site DNA-binding activity and an NF-κB protein that appears mostly processed and in the nucleus of cells. NF-κB DNA-binding activity and processing is increased by treatment of sponge tissue with LPS. By transcriptomic analysis of *A. queenslandica* we identified likely homologs to many upstream NF-κB pathway components. These results present a functional characterization of the most ancient metazoan NF-κB protein to date, and show that many characteristics of mammalian NF-κB are conserved in sponge NF-κB, but the mechanism by which NF-κB functions and is regulated in the sponge may be somewhat different.

---

Sponges are the sole members of the phylum Porifera (1), with about 8,500 known extant species (1). They are widely considered the most basal metazoans (2) and the sister group to all other multicellular animals (3). Sponges are estimated to have evolved prior to the Cambrian explosion, probably appearing approximately 600 million years ago (4). Sponges are found in a variety of aquatic habitats throughout the world, with the majority of species living in marine environments (1).

Sponges have simple body plans that resemble a sack of cells with an extracellular matrix and pores. Nevertheless, the simplicity of the sponge body plan belies the complex mechanisms that must be coordinated to allow for their existence. Indeed, due to their water filtration system and body plan, sponges have one of the highest exposures to a diverse array of microorganisms as compared to other multicellular organisms (5). Therefore, it is not surprising that they have a robust innate immune system to monitor and counteract this continuous exposure to microbes. As such, sponges have the ability to distinguish between friendly and foreign matter, such as the recognition of cells from heterologous sponges, the uptake of bacteria by phagocytosis through macrophage-like cells (6), the production of secondary metabolites for defense (7), and the recognition of bacterial cell wall-derived lipopolysaccharide (LPS) (8). The molecular mechanisms by which these defense processes occur are largely unknown; however, the presence of homologs to higher metazoan innate immune molecules in sponges suggests a complex level of immune defense and regulation (9).

Because of their phylogenetic position at the earliest lineage of living animals, sponges are useful for addressing questions regarding the origin of metazoan-specific gene families and their evolution (10). The Great Barrier Reef demosponge *Amphimedon queenslandica* (Aq) is the most extensively studied poriferan, including having its genome sequenced and being the subject of several studies, mostly focused on investigating the evolution of metazoan development and multicellularity (11–14).

One of the most prominent pathways involved in the regulation of immunity in advanced metazoans is transcription factor NF-κB, which is regulated by many upstream factors (15). Many homologs of the NF-κB pathway are conserved across diverse phyla, from vertebrates to the single-celled holozoan *Capsaspora owczarzaki* (16). NF-κB pathway components and transcriptional responses have been intensively studied in vertebrates, and to some extent flies, for their involvement in immunity, development, and disease. However, this pathway and genes regulated by NF-κB are, for the most part, less well-characterized in more evolutionary basal organisms (16).

Proteins in the NF-κB superfamily are related through the Rel Homology Domain (RHD), which contains sequences important for dimerization, DNA binding, and nuclear translocation (15). The two subfamilies of NF-κB proteins comprise the traditional NF-κBs (p52/p100, p50/p105, *Drosophila* Relish), and the Rel proteins (RelA, RelB, c-Rel, *Drosophila* Dif and Dorsal). Rel and NF-κB proteins differ in their DNA-binding preferences as well as their C-terminal domain structures. For example, NF-κB proteins contain C-terminal inhibitory sequences called Ankyrin (ANK) repeats, whereas Rel proteins contain C-terminal transactivation domains. Overall, the amino acid (aa) sequence and structural organization of NF-κB proteins in cnidarians (16–19) poriferans (20), the protist *Capsaspora owczarzaki* (21), and some choanoflagellates (22) more closely resemble the NF-κB subfamily of proteins found in vertebrates.

Activation of NF-κB pathways can be initiated by various upstream ligands that bind to their specific receptors (e.g., Toll-like receptors (TLRs), Interluekin-1 receptors (IL-1Rs), tumor necrosis factor receptors (TNFRs)), which direct appropriate downstream effects. For activation of the human non-canonical pathway, NF-κB p100 undergoes an IKK-mediated phosphorylation on a cluster of serine residues located C-terminal to the ANK repeats. p100 is then ubiquitinated and C-terminal sequences are selectively degraded by the proteasome to release the active NF-κB p52 protein (23). In some cases in humans, Tank-binding Kinase (TBK) functions with other signaling complexes that feed into this NIK-IKK cascade, which leads to NF-κB activation (24). Non-canonical processing of p100 ultimately leads to generation of the RHD-only protein p52, which can then translocate to the nucleus, bind DNA, and activate target genes (15).

Transcriptomic and genomic sequencing has revealed that NF-κB and homologs of many of its upstream regulators are present in most eukaryotes from protists to vertebrates (16). However, the numbers and structures of these signaling proteins vary across species, and generally become more complex and numerous through evolutionary time.

In this manuscript, we have used phylogenetic, biochemical, and cell- and tissue-based assays to characterize the structure, activity, and regulation of sponge NF-κB. Our results indicate that sponge NF-κB has many of the same properties as other metazoan NF-κB proteins, but also has sequences and methods of regulation that may be specific to poriferans.

## Results

### A. queenslandica *NF-κB resembles other metazoan NF-κB proteins both structurally and phylogenetically*

Gauthier & Degnan (2008) reported that the sole NF-κB protein in the demosponge sole *A. queenslandica* (Aq-NF-κB) has an overall structure (RHD-ANK repeat domain) and sequence phylogeny that are more closely related to mammalian NF-κB proteins (p100/105) than Rel proteins (RelA, RelB, c-Rel) proteins. That is, Aq-NF-κB contains an N-terminal RHD, a nuclear localization sequence (NLS), a glycine-rich region (GRR), and ANK repeats (Fig. 1A). However, the Aq-NF-κB has an additional 171-aa sequence between the GRR and the ANK repeats (Fig. 1A, grey domain) that is not found in any other NF-κB protein and has no sequence similarity to any other protein in BLAST databases. Nevertheless, the general structure of Aq-NF-κB resembles other basal NF-κBs discovered to date, such as those in cnidarians (17–19) and *Capsaspora owczarzaki* (21), as well as the human p100 and p105 proteins (Fig. 1B). Our own phylogenetic comparison of RHD sequences confirmed that the Aq-RHD is more similar to the RHDs of other NF-κB proteins as compared to Rel proteins across phyla, and that the Aq-RHD sequences are most closely related to NF-κB-like proteins Cnidaria, the closest metazoan phylum to Porifera (Fig. 1C). Taken together, these results are consistent with other studies indicating that NF-κB proteins are more ancient than Rel proteins (16–19).

**Figure 1.**
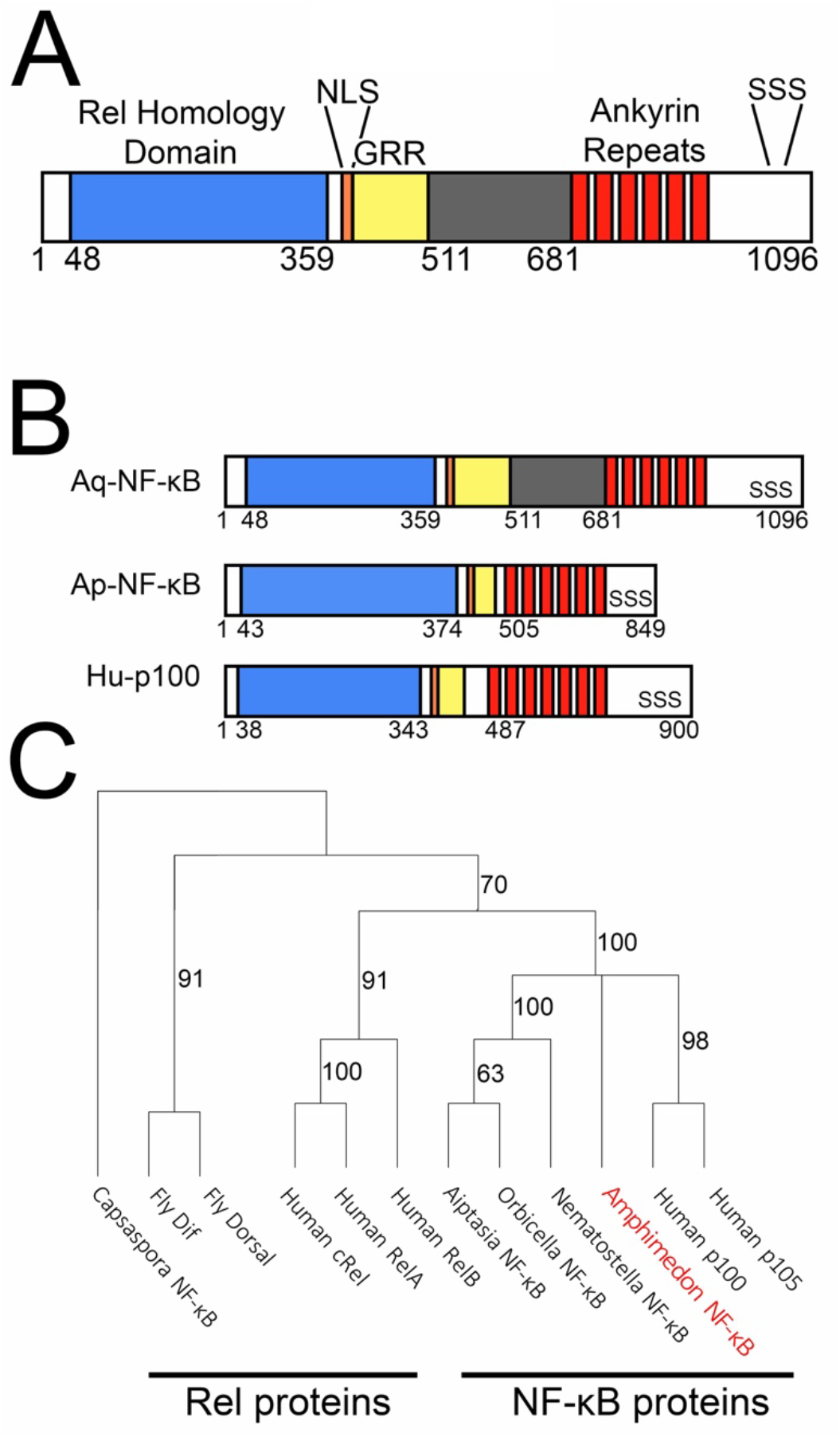
The *A. queenslandica* NF-κB protein resembles other metazoan NF-κB proteins both structurally and phylogenetically. *A*, Schematic of Aq-NF-κB domain structures. Rel Homology Domain (RHD, blue), Nuclear Localization Sequence (NLS, orange), Glycine Rich Region (GRR, yellow), extra sequences after GRR (grey), Ankryin Repeats (ANK, red), and conserved C-terminal serines (SSS). *B*, Schematic comparing domain structures of Aq-NF-κB (1096 amino acids), the previously characterized sea anemone Aiptasia NF-κB protein (Ap-NF-κB), and human p100. Functional domains shared between these proteins are marked by colors, as in A. *C*, Phylogenetic analysis of RHD sequences of the indicated NF-κB proteins was performed using distance neighbor-joining tree analysis. The phylogeny was rooted with the predicted RHD of the single-celled protist *Capsaspora owczarzaki*. Branches indicate bootstrap support values.

### Aq-NF-κB protein contains sequences that bind DNA and activate transcription

To investigate the overall DNA binding-site specificity of Aq-NF-κB, we assessed the DNA-binding profile of bacterially expressed, affinity purified RHD sequences of Aq-NF-κB in a protein binding microarray (PBM) consisting of 2592 κB sites and 1159 random background sequences (for sequences see Mansfield et al., 2017). We (17,25) and others (26) have previously used this type of analysis on both mammalian and cnidarian NF-κB proteins. By analysis of z-scores for binding sites on the PBM, the DNA-binding profile of Aq-NF-κB is similar to NF-κB from the sea anemone *N. vectensis* and human NF-κB p52, but is distinct from human c-Rel and human RelA (Fig. 2A).

**Figure 2.**
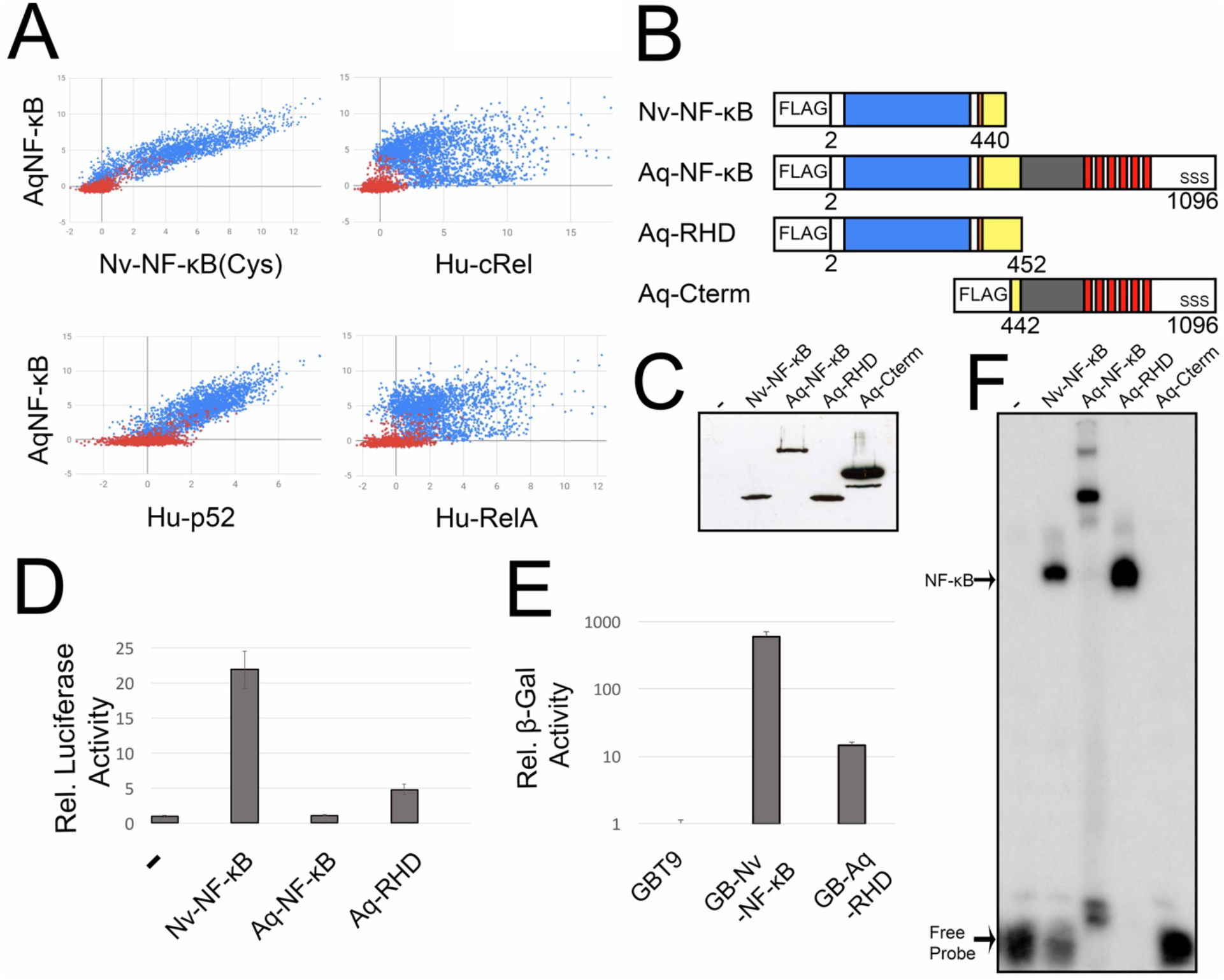
Aq-NF-κB protein contains sequences that bind DNA are transcriptionally active. *A*, DNA-binding profiles of Aq-NF-κB in a protein binding microarray (PBM) with *Nematostella* (Nv) NF-κB (cysteine (cys) allele) (top, left), Human cRel (top, right), p52 (bottom, left), and RelA (bottom, right). Red dots represent random background sequences and blue dots represent κB binding sites. *B*, FLAG-tagged expression vectors used in these experiments. From top to bottom the drawings depict the naturally shortened Nv-NF-κB, the full-length Aq-NF-κB protein, an N-terminal-only mutant containing the RHD and glycine rich region (Aq-RHD), and a C-terminal-only mutant containing the ANK repeats and other C-terminal sequences (Aq-Cterm). *C*, Anti-FLAG Western blot of 293T cell lysates transfected with the indicated expression vectors. *D*, A κB-site luciferase reporter gene assay was performed with the indicated proteins in 293 cells. Luciferase activity is relative (Rel.) to that seen with the empty vector control (1.0), and values are averages of three assays performed with triplicate samples with standard error. *E*, A GAL4-site *LacZ* reporter gene assay was performed in yeast Y190 cells. β-galactosidase (β-gal) reporter gene activity is relative (Rel.) to the GAL4 (aa 1-147) control (1.0) and is presented on a log-scale. Values are averages of seven assays performed with duplicate samples with standard error. *F*, A κB-site electromobility shift assay (EMSA) using each of the indicated lysates from C. The NF-κB complex and the free probes are indicated by arrows.

In mammals and cnidarians, NF-κB proteins require post-translational processing for nuclear translocation, active DNA binding, and transcriptional activation activity (17,18,23). To investigate corresponding properties of Aq-NF-κB, we first created FLAG-tagged expression vectors for the full-length Aq-NF-κB, a mutant containing the RHD and GRR, and a mutant containing only the sequences C-terminal to the RHD (Aq-Cterm, i.e., primarily the ANK repeats) (Fig. 2B). As a control, we used an expression vector for the *N. vectensis* NF-κB protein (Nv-NF-κB), which we have characterized previously as an active, naturally truncated NF-κB protein (17–19). As shown by anti-FLAG Western blotting (Fig. 2C), each construct expressed a protein of the appropriate size when transfected into HEK 293T cells.

To analyze the transactivation properties of the full-length and the truncated Aq-NF-κB proteins, we performed an NF-κB-site luciferase reporter assay in 293 cells. Aq-RHD and Nv-NF-κB both activated transcription of the reporter gene (4.8-fold and 21.9-fold, respectively) as compared to the empty vector control and to full-length Aq-NF-κB (Fig. 2D). Furthermore, a GAL4-Aq-NF-κB RHD fusion protein activated transcription approximately 15-fold above control levels in a GAL4-site reporter assay in yeast (Fig. 2E). As we have shown previously (18,19,27), a GAL4-Nv-NF-κB fusion protein strongly activated transcription in yeast (approximately 600-fold) as compared to the GAL4 alone control. Therefore, the ability of the Aq-RHD sequences to activate transcription appears to be an intrinsic property of sequences within the N-terminal half of the protein, and this transactivaton activity is not apparent in the full-length Aq-NF-κB protein.

To assess the DNA-binding ability of Aq-NF-κB, whole-cell detergent extracts from 293T cells transfected with the Aq-NF-κB expression plasmids were analyzed in an electrophoretic mobility shift assay (EMSA) using a κB-site probe, which we have previously shown can be avidly bound by cnidarian NF-κB proteins (17–19). Lysates from cells transfected with expression plasmids for Aq-RHD and the positive control Nv-NF-κB both contained an activity that strongly bound to the κB-site probe (Fig. 2F). As expected, lysates from cells transfected with the empty vector or the Aq-Cterm did not have DNA-binding activity. Somewhat surprisingly, full-length Aq-NF-κB bound the κB-site probe to nearly the same extent as the Aq-RHD protein. Taken together, these results demonstrate that removal of C-terminal ANK repeat domain sequences of Aq-NF-κB enables the protein to activate transcription in vertebrate cells, but the presence of C-terminal sequences of Aq-NF-κB does not substantially inhibit its DNA-binding activity when Aq-NF-κB is expressed in human cells.

### C-terminal truncation of Aq-NF-κB enables it to translocate to the nucleus and can be induced by an IKK-dependent mechanism in vertebrate cells

In non-canonical NF-κB pathway activation, the human NF-κB p100 protein is converted to an active form by proteasomal processing of C-terminal sequences up to the GRR, which allows for the shortened p52 protein to translocate to the nucleus (23). To investigate the subcellular localization properties of Aq-NF-κB, we expressed the above described FLAG-tagged Aq-NF-κB proteins in DF-1 chicken fibroblast cells and assessed their subcellular localization by anti-FLAG indirect immunofluorescence (Fig. 3A). Full-length Aq-NF-κB showed cytoplasmic staining, whereas the C-terminally truncated Aq-RHD protein was located exclusively in the nucleus (as judged by overlap with DAPI staining of the nucleus). The N-terminally deleted Aq-Cterm protein, which consists of essentially the C-terminal ANK repeats, showed cytoplasmic staining. Consistent with our previous results (19), the naturally shortened *N. vectensis* Nv-NF-κB protein localized to the nucleus in these cells.

**Figure 3.**
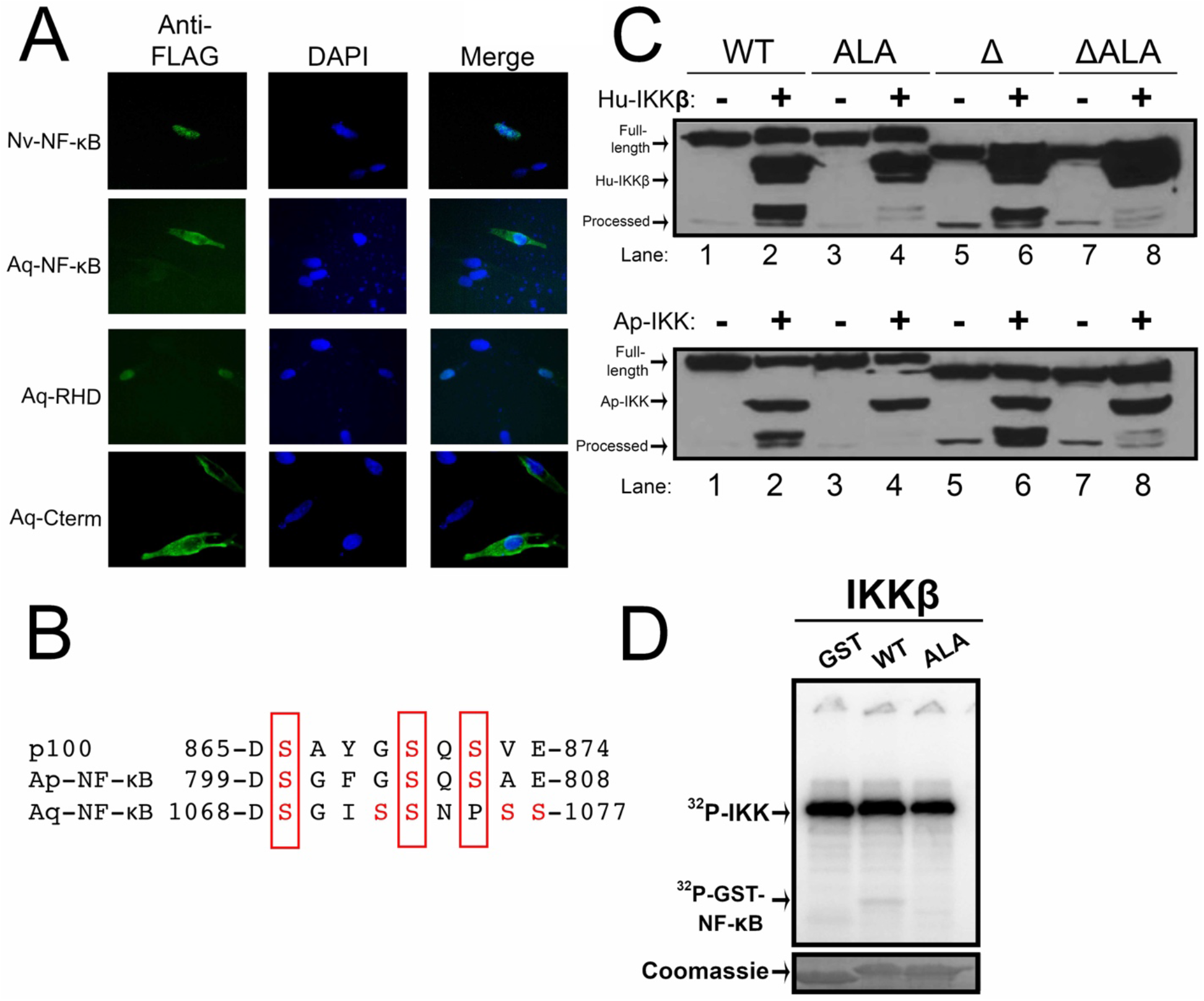
Analysis of C-terminal sequences of Aq-NF-κB for inhibition of nuclear translocation and for IKK-dependent phosphorylation and induced degradation. *A*, DF-1 chicken fibroblasts were transfected with the indicated FLAG vectors and subjected to anti-FLAG indirect immunofluorescence. Anti-FLAG, green (left); DAPI-stained nuclei, blue (middle); and merged images (right). *B*, An alignment of relevant regions of NF-κB from human p100, the sea anemone Aiptasia, and *A. queenslandica*. Conserved serine residues are indicated in red. *C*, Western blot of lysates from 293T cells co-transfected with Aq-NF-κB wild-type (WT), C-terminal alanine mutant (ALA), N-terminal deletion mutant (Δ), or double mutant (ΔALA) and with empty vector (−) or FLAG-Hu-IKKβ (top) or FLAG-Ap-IKK (bottom). Full-length and processed forms of Aq-NF-κB, and FLAG-Hu-IKKβ are indicated. Lanes are numbered at the bottom. *D, In vitro* kinase assay (top) using FLAG-Hu-IKKβ protein with bacterially expressed substrates of GST alone or GST fusion proteins containing the wild-type (WT) serine residues of Aq-NF-κB (amino acids 1048-1086), or the same C-terminal residues with conserved serines mutated to alanines (ALA). The GST proteins were electrophoresed on an SDS-polyacrylamide gel and stained with Coomassie blue (bottom).

We have previously shown that two cnidarian NF-κB proteins (from the sea anemone Aiptasia and the coral *Orbicella faveolata*) can be induced to undergo C-terminal processing in human 293T cells by overexpression of human and cnidarian IKKs (17,18). Furthermore, IKK-induced processing of these cnidarian NF-κBs requires a cluster of C-terminal serine residues with similar sequence and spacing to ones found in human NF-κB p100 (17,18,23). Previously, results from our lab indicated that co-transfection of IKKs with Aq-NF-κB did not its induce C-terminal processing (17). This lack of processing was proposed to be due to the absence of obvious sequence homology of sites of IKK phosphorylation in the C-terminal sequences of Aq-NF-κB as compared to human p100 and Aiptasia NF-κB. Based on that result, we hypothesized that the native spacing of serines near the C-terminus of Aq-NF-κB was not allowing IKK-induced C-terminal processing of the protein. Therefore, in initial experiments, we created a FLAG-tagged Aq-NF-κB mutant that had the same spacing and number of serines as Aiptasia NF-κB (Supplemental Fig. 1A). WT Aq-NF-κB and the Aq-NF-κB mutant with the Aiptasia-like serine cluster were then cotransfected with empty vector, Hu-IKKβ, or Ap-IKK. In these experiments, WT Aq-NF-κB and the Aq-NF-κB-Aiptasia serine mutant were both processed when transfected with the IKKs, but neither was processed in cells transfected with the empty vector control (Fig. 3C, top and bottom, lanes 1 and 2, and Supplemental Fig. 1B). This result demonstrated that the WT Aq-NF-κB can indeed be processed in 293T cells by co-expression of these IKKs, and that there was likely an error in analysis in these previous (17) experiments. We believe that the previously published Western blot was run too long and the lower processed band was run off the gel.

With this new information, we sought to identify sequences important for IKK-induced C-terminal processing of Aq-NF-κB. In a reanalysis, we found that Aq-NF-κB contains a cluster of serine residues that are somewhat similar to the serines required for processing of human p100 and the cnidarian NF-κB proteins (Fig. 3B, serines in red). Additionally, the initial discovery of Aq-NF-κB identified extra sequences after the GRR (aa 511-681) (20) that contained several serines, but the function of this region is not known (Fig. 1A). To determine which sequences were required for processing of Aq-NF-κB, we created three mutants: 1) with five C-terminal serine residues (Fig. 3B serines in red) changed to phospho-resistant alanines (ALA), 2) with a deletion of the additional region between the GRR and the ANK repeats (Δ), and 3) a double mutant (ΔALA) that had the serine-to-alanine mutations and the deletion of aa 511-681 region. The Aq-NF-κB-ΔLA showed substantially reduced IKK-induced processing as compared to wild-type Aq-NF-κB in 293T cells (Fig. 3C, top and bottom, lanes 3 and 4). The Aq-NF-κBΔ mutant showed IKK-induced processing similar to WT Aq-NF-κB (Fig. 3C, top and bottom, lanes 5 and 6), and the double mutant Aq-NF-κB-ΔALA showed reduced processing as compared to the Aq-NF-κBΔ mutant (Fig. 3C, top and bottom, lanes 7 and 8). Overall, these results demonstrate that IKK-induced C-terminal processing can also occur with Aq-NF-κB (17–19), and that this processing requires a cluster of C-terminal serine residues but does not require the additional sequences (aa 511-681) in Aq-NF-κB.

To determine whether these serine residues could be directly phosphorylated by human IKKβ, we first generated bacterially expressed GST-fusion peptides containing either the wild-type or phospho-resistant alanine sequences (aa 1048-1056) near the C-terminus of the Aq-NF-κB. These GST-fusion proteins were then incubated with FLAG-Hu-IKKβ in a radioactive *in vitro* kinase assay. As shown in Fig. 3D, the GST-tagged peptides containing the wild-type Aq-NF-κB serine residues, but not the Ala residues, were phosphorylated by IKKβ.

### Sponge tissue expresses putative NF-κB proteins and contains NF-κB site DNA-binding activity that increases following treatment with bacterial lipopolysaccharide (LPS)

To determine whether sponge tissue expresses active NF-κB, we acquired tissue from an aquarium grown black encrusting demosponge from the Caribbean that is likely of the genus *Cliona* (Fig. 4A). This sponge was chosen because of the difficulty of growing *A. queenslandica* in captivity (28) and because preliminary experiments showed that our anti-sea anemone NF-κB antiserum (17) cross-reacted with our aquarium sponge NF-κB protein but not with Aq-NF-κB (not shown). Anti-NF-κB Western blotting of extracts from the black encrusting sponge tissue detected one high (∼130 kDa) and one lower (∼65 kDa) molecular weight band, with the majority of the protein being in the lower band (Fig. 4B). As expected, the anti-Ap-NF-κB antibody detected two bands of appropriate sizes in a 293T cell extract containing overexpressed Aiptaisa NF-κB (Fig. 4B).

**Figure 4.**
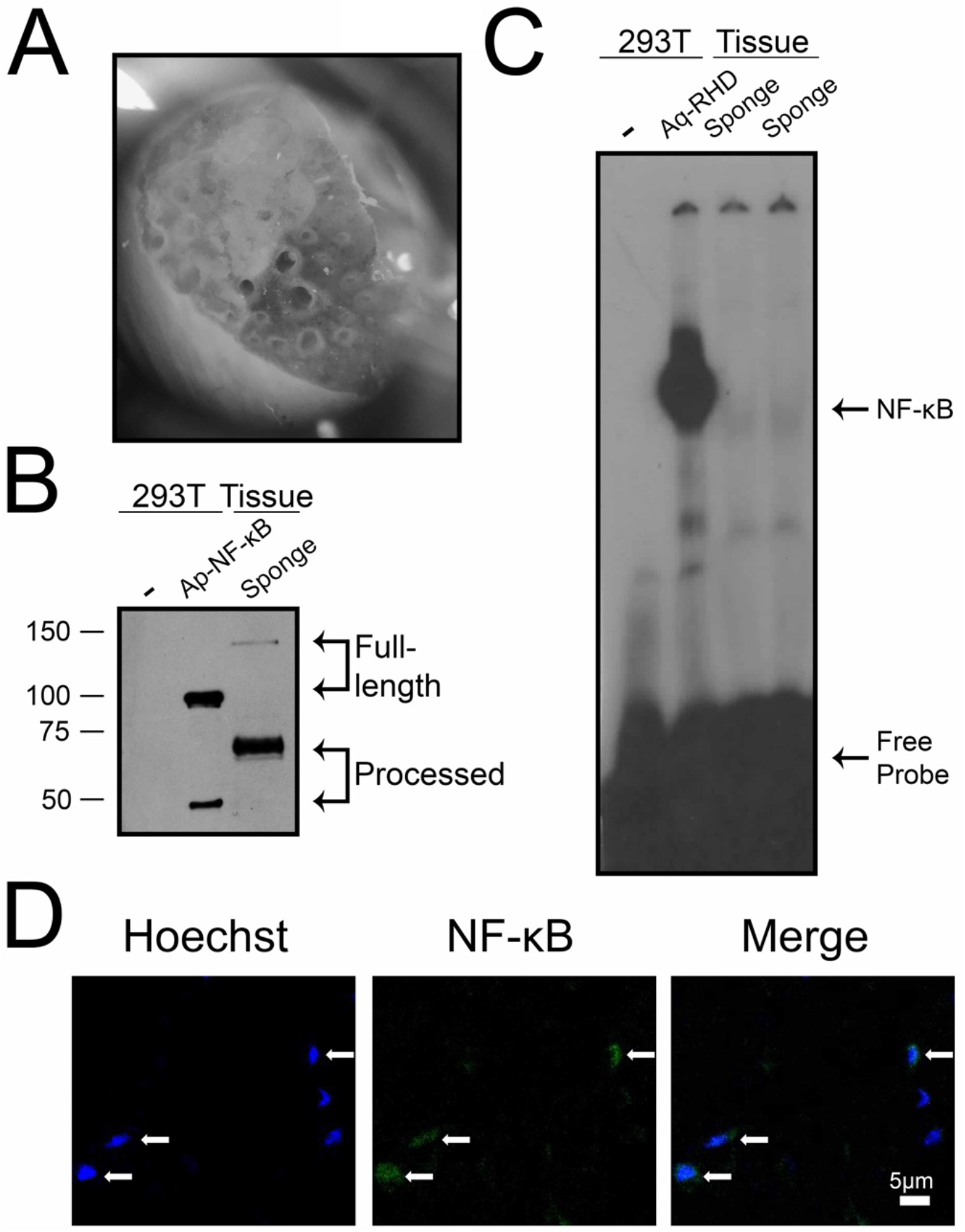
Sponge tissue extract contains NF-κB-like proteins that have DNA-binding activity and localize to the nucleus. *A*, Black encrusting sponge tissue under a light microscope. *B*, Anti-Ap-NF-κB Western blot of lysates from 293 cell lysates transfected with either empty vector (first lane) or FLAG-Ap-NF-κB (middle lane), and a lysate from black encrusting sponge tissue (third lane). Molecular weight markers are indicated to the left of the blot in kilodaltons. *C*, A κB-site EMSA with 293 cell lysates transfected with either empty vector (first lane) or FLAG-Aq-RHD (second lane), or with tissue lysates from black encrusting sponge tissue (third and fourth lanes). NF-κB complex shift and free probe are indicated by arrows. *D*, Confocal microscopy of cryosectioned tissue demonstrates some cells in sponge tissue express NF-κB, and most of the expression is localized to the nucleus. Hoechst-stained nuclei are shown in blue (left panel), NF-κB is in green (middle panel), and a merged image showing co-localization (right panel).

When this same sponge tissue lysate was analyzed by an EMSA using an NF-κB-site probe, a band was detected that migrated similar to the prominent band seen with extracts from 293T cells overexpressing FLAG-Aq-RHD (Fig. 4C).

To analyze the organismal pattern of NF-κB expression, we performed anti-Ap-NF-κB immunofluorescence on cryosections of black encrusting sponge tissue. A subset of cells expressed immunoreactive proteins that closely coincided with Hoechst-stained nuclei in the tissue (Fig. 4D).

Overall, the results in this section indicate that the black encrusting sponge tissue has proteins that are similar to full-length and processed NF-κB proteins from other species, contains NF-κB-site binding activity, and expresses NF-κB in the nucleus of some but not all cells.

To determine whether a known activator of mammalian NF-κB could affect sponge NF-κB, we treated black encrusting sponge tissue with *E. coli* lipopolysaccharide (LPS) for 30 min and then analyzed the tissue by Western blotting or EMSA. After treatment with LPS, the amount of processed protein (lower band, Fig. 5A) increased by approximately 15% and the amount of unprocessed protein decreased by approximately 10% (normalized to the α-tubulin loading control, Supplemental Fig. 2). Additionally, LPS treatment resulted in increased κB-site DNA-binding activity by EMSA (Fig. 5B). Overall these results demonstrate that treatment of the isolated sponge tissue with LPS results in an increase in NF-κB processing and DNA-binding activity.

**Figure 5.**
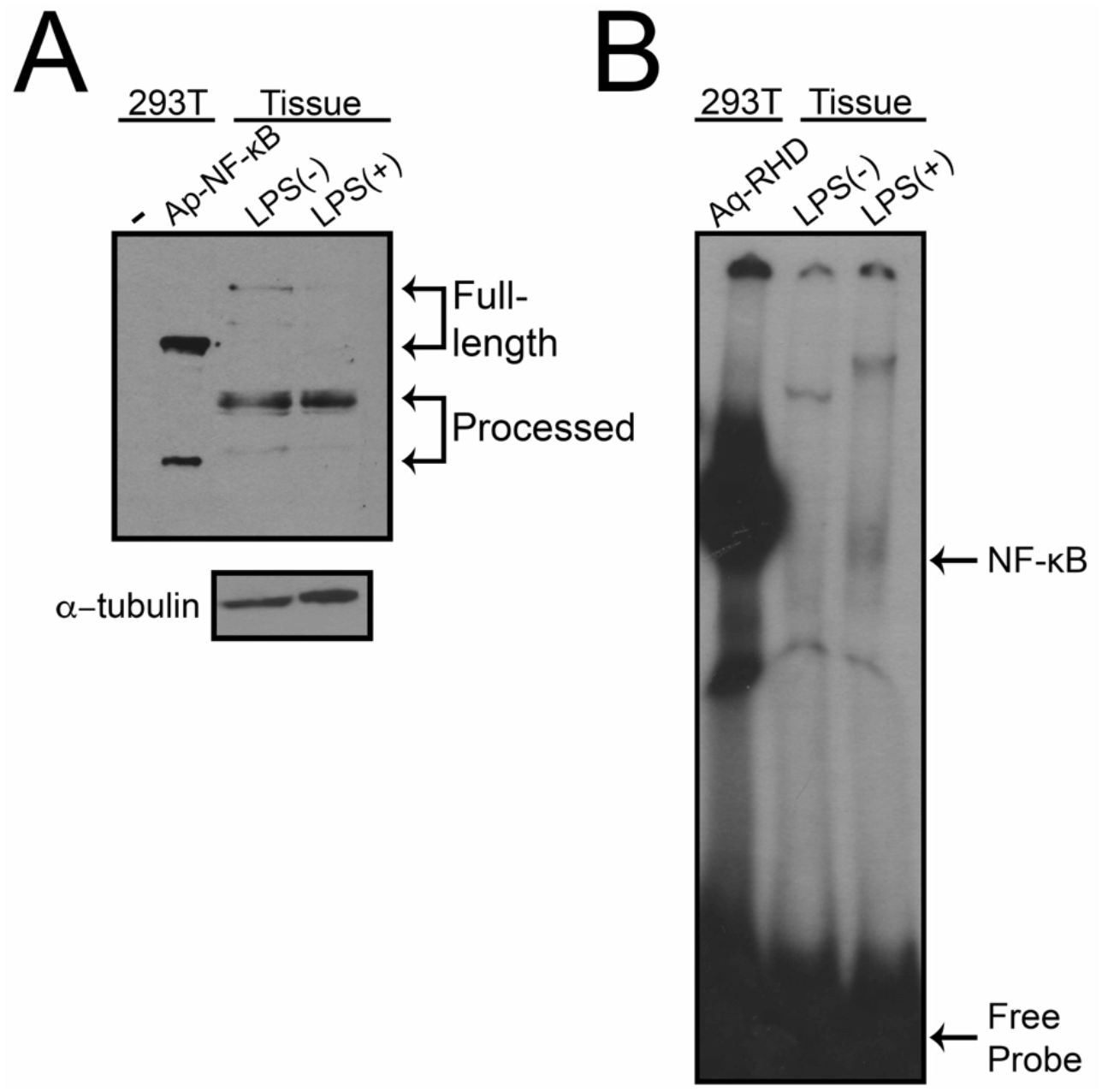
Treatment of sponge tissue with LPS results in increased NF-κB protein and activity. Black encrusting sponge tissue was treated with control (−) or lipopolysaccharide (LPS) conditions for 30 min. Lysates were then prepared and analyzed by anti-NF-κB Western blotting (*A*) or EMSA (*B). A*, Lanes one and two contain 293T cell extracts transfected with empty vector or FLAG-Ap-NF-κB, respectively. Lanes three and four contain black encrusting sponge tissue lysates either untreated (lane three) or treated (lane four) with LPS. α-tubulin Western blotting is shown below for normalization purposes. *B*, A κB-site EMSA with 293 cell lysates from FLAG-Aq-RHD (first lane), or tissue lysates from untreated or LPS-treated black encrusting sponge tissue (third and fourth lanes, respectively). The NF-κB-DNA complexes and the free probes are indicated.

## Discussion

In this report, we demonstrate that at least two sponges have active NF-κB proteins. That is, we show that the *A. queenslandica* NF-κB protein is most phylogenetically and structurally similar to other NF-κB proteins (Fig. 1), and that its DNA-binding profile, transactivation properties and regulation by C-terminal sequences are also similar to NF-κB proteins from other species (Fig. 2). We also demonstrate that a black encrusting demosponge contains proteins that can bind a κB site in an EMSA and are detected by an anti-NF-κB antibody in Western blots and tissue immunofluorescence (Fig. 4). Furthermore, treatment of this sponge tissue with LPS leads to further activation of its putative NF-κB protein, as shown by both increased processing and increased DNA binding (Fig. 5). Finally, we also show that *A. queenslandica* contains homologs to many components of a TLR-to-NF-κB pathway (Fig. 6 and Table 1).

**Figure 6.**
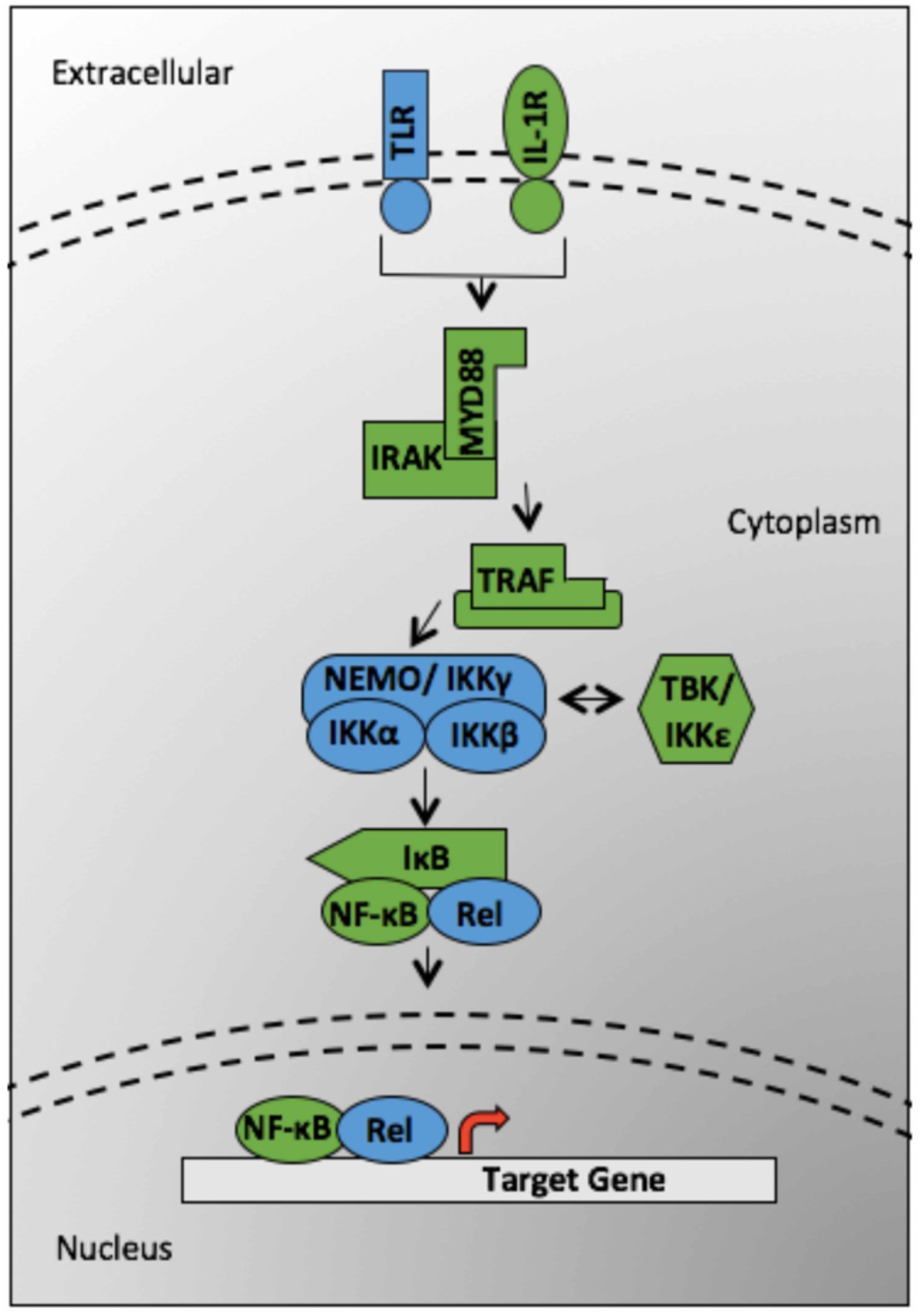
Transcriptomic analysis reveals homologs of NF-κB pathway and upstream signal transduction receptors. NF-κB pathway members that are present in Homo sapiens are colored in blue, and homologs that are present in the *A. queenslandica* genome are colored in green.

**Table 1.** TLR-to-NF-κB pathway homologs encoded in the *A. queenslandica* genome. Listed are selected proteins in the pathway and their Accession numbers on NCBI. “Blast Accession Number” indicates the protein sequence used to identify the *A. queenslandica* homolog. Comments provide details of domain structures of these various sequences.

| Protein Name | Blast Accession Number | Homolog Present *A. queenslandica* | Accession Number of *A. queenslandica* Homolog | Homologs in *A. queenslandica* Homolog | Comments |
| --- | --- | --- | --- | --- | --- |
| TLR | NP_003254.2 | - |  |  |  |
| IL-1R |  | + | XP_003383414.2 | 1 | TIR-Only protein with Immunoglobulin-like domains |
| MYD88 | NP_002459.3 | + | NP_001266224.1 | 1 |  |
| IRAK | NP_066961.2 | + | XP_019855709.1<br>XP_019855710.1 | 2 | Only has Kinase domain (missing death domains) |
| TRAF | NP_066961.2 | + | XP_019855421.1<br>XP_019861351.1<br>XP_003383545.3<br>XP_019858615.1<br>XP_019858616.1<br>XP_019858613.1<br>XP_003386185.1<br>XP_003387035.1<br>XP_019852875.1 | 13 | Most have ring domains and MATH domains, while some have Zn-coil domains and Zn-finger domains |
|  |  |  | XP_019852874.1<br>XP_003385751.1<br>XP_019860776.1<br>XP_019852876.1 |  |  |
| NEMO/IKKγ | NP_003630.1 | - | | | |
| IKKα | NP_001269.3 | - | | | |
| IKKβ | NP_001547.1 | - | | | |
| TBK/IKKε | NP_037386.1 | + | XP_019854889.1<br>XP_003387126.1 | 2 | |
| IκB | NP_113607.1<br>NP_004547.2<br>NP_002494.2 | + | XP_011404515.1<br>XP_019853663.1<br>XP_019853661.1 | 3 | Epsilon-like<br>Cactus-like |
| NF-κB | NP_001070962<br>.1 | + | NP_001266238.1 | 1 | |
| Rel | NP_006500.2 | - |  |  |  |

Consistent with previous results (20), our results indicate that Aq-NF-κB is structurally similar to the human NF-κB p100 protein and to our previously characterized sea anemone *Aiptasia* and coral *O. faveolata* NF-κB proteins (17,18,23). In our cell-based assays, removal of the ANK repeat domain allowed for localization of Aq-NF-κB to the nucleus and liberation of its transcription activation activity.

The occurrence of bipartite NF-κB proteins that contain both the RHD and the C-terminal ANK repeats in sponges (A. *queenslandica*) and the protist *C. owczarzaki* is different than what is seen in some cnidarians, such as *N. vectensis*, that contain separate RHD and IκB-like ANK-repeat proteins. More specifically, we and others have demonstrated that the sponge and protist NF-κB proteins in metazoans are more closely related structurally to the human p100 and p105 NF-κB proteins (17,18,20), suggesting that genes such as the naturally shortened Nv-NF-κB protein (19) are the result of organism-specific gene splitting events. Such gene-splitting events are not uncommon in cnidarians (16).

Surprisingly, full-length Aq-NF-κB bound to DNA to a similar extent as the Aq-RHD protein (deleted of the C-terminal ANK repeats) (Fig. 2F). Therefore, the C-terminal ANK repeat domain of Aq-NF-κB appears to inhibit nuclear translocation (Fig. 3A), but not DNA binding. In all other NF-κB proteins analyzed to date, the C-terminal ANK repeat domain inhibits the DNA-binding activity of RHD sequences in assays similar to those that we have performed here. As we note, the Aq-NF-κB protein contains an unusually long sequence between its RHD and the start of its ANK repeats. It is possible that this extended sequence interferes with the ability of the ANK repeat domain to inhibit DNA binding by the RHD sequences. In any case, the ANK repeat domain can block nuclear translocation (Fig. 3A) and transcriptional activation by Aq-NF-κB in reporter gene assays (Fig. 2D), and inhibition of nuclear translocation would functionally block Aq-NF-κB’s ability to activate gene transcription in cells. Why there are extra sequences between the RHD and the ANK repeat domain in Aq-NF-κB and whether they are important for any NF-κB–related processes in the sponge are not known. However, the additional sequences do not appear to be involved in or required for proteolytic processing of Aq-NF-κB by known IKKs (Fig. 3C).

*A. queenslandica* is similar to other evolutionarily basal organisms in that its genome encodes one NF-κB protein and no Rel proteins. Additionally, Aq-NF-κB contains serine residues C-terminal to the ANK repeats that can be phosphorylated *in vitro* by a human IKK protein, and these conserved serines are required for IKK-induced processing of Aq-NF-κB in human cells in tissue culture. Nevertheless, the majority of a putative NF-κB protein in tissue from a black encrusting sponge appears to be processed, binds to DNA, and is located in the nucleus of cells. Thus, in this sponge, most of its NF-κB protein appears to be constitutively in its active form. We have previously shown that most of the NF-κB protein of the sea anemone Aiptasia is also a constitutively nuclear and processed protein in animals (17). Moreover, the majority of mouse NF-κB p100 is in its processed form when analyzed in extracts taken directly from liver, lymph nodes, and bone marrow (29). What factors promote constitutive processing of NF-κB proteins in these diverse organisms and tissues *in vivo* is not known. However, it is interesting to speculate that continual exposure to pathogens or other activating ligands induces NF-κB processing in each of these situations. Further studies should be directed at determining the requirements for the production of the constitutively active sponge NF-κB protein, whether this constitutive activity is also found in other poriferans, the trade-offs to this constitutive activity, and the situations in which this processing does not occur.

Two IKK-like proteins have been identified in genomic sequences of *A. queenslandica* (NCBI Accession numbers XP_019854889.1 and XP_003387126.1), and both appear more similar to the TBK/IKKε subclass of IKKs than to the IKKα/β subclass (data not shown). In general, the mammalian TBK/IKKε kinases do not phosphorylate IκBs or C-terminal serines in p100. Therefore, although our results in 293T cells suggest that *A. queenslandica* has a kinase that is capable of inducing processing of Aq-NF-κB, such a kinase has yet to be identified, either because there is a lack of genomic coverage to identify it or there is an absence of obvious sequence homology to other known metazoan IKKs.

Although we have yet to identify the kinase capable of processing Aq-NF-κB, we have identified many other homologs of a likely NF-κB pathway in *A. queenslandica* (Fig. 6). *A. queenslandica* lacks a complete Toll-like receptor (TLR), but does expresses what appears to be a homolog of the Interleukin 1 receptor (IL-1R), which contains the intracellular Toll-interleukin receptor (TIR) domain as well as the extracellular immunoglobulin domains. While TLR and IL-1R bind different ligands due to their different extracellular domains, both receptors contain intracellular TIR domains which can interact with many of the same downstream signaling proteins.

We also found that treatment of fresh sponge tissue with LPS for 30 min resulted in the almost complete processing of full-length NF-κB protein, and an increase in κB-site binding activity. In the demosponge *Suberites domuncula* it has been shown that LPS can be bound by an LPS-interacting protein (5), which is capable of binding and dimerizing with an Sd-MYD88 homologous protein. Currently, there is no identified LPS-interacting protein homolog in the *A. queenslandica*, but it does contain a homolog of the TIR-domain adapter protein MYD88 (Table 1). However, our data suggest that the black encrusting sponge possesses an LPS-interacting protein or a receptor capable of recognizing LPS.

Recently, it was shown that *A. queenslandica* NF-κB transcripts are expressed in at least two cell types: choanocytes, which are cells that reside in the chambers that allow for the sponge to filter feed, and to a lesser extent in archaeocytes, which are pluripotent cells that are motile inside the mesoglea (30). In the black encrusting sponge that we examined, primarily nuclear NF-κB protein was detected in a subset of cells that appeared to be within the tissue mesoglea. We did not detect any choanocyte chambers in our images. Therefore, we believe that the cells that are stained with our NF-κB antiserum are likely either archaeocytes or another cell type that has yet to be identified. Archaeocytes are known to be amoeboid-like and to transport particulate matter taken-up by choanocytes into the sponge tissue, so it is possible that they require NF-κB-related immune functions. Furthermore, archaeocytes are also known to be totipotent and capable of differentiation into many cell types (31), and it is well established that NF-κB plays a role in differentiation of different cell types in vertebrates (32–34).

The ability of LPS to further activate NF-κB in sponge tissue suggests that NF-κB has role in poriferan anti-bacterial immunity. However, there is likely to also be an early developmental role for NF-κB in poriferans. That is, Gauthier & Degnan (2008) showed that NF-κB transcripts are present during early development of *A. queenslandica*, especially in granular cells and flask cells (20). Such dual roles in adult immunity and early development have been shown for the NF-κB protein Dorsal in *Drosophila* and has been postulated for the NF-κB protein in *N. vectensis* (19).

Although the last common ancestor of sponges and *Homo sapiens* was approximately 600 million years ago, the structural and functional conservation demonstrated here suggests that regulated processing of NF-κB is an ancient property. Our data provide a functional analysis of the most basal NF-κB protein to date, and provide the first experimental connection of NF-κB to immunity in our oldest animal ancestor, the sponge.

## Experimental procedures

### Phylogenetic analyses

The RHD sequences of NF-κB from *A. queenslandica* (Aq) (20) were compared phylogenetically to *Aiptasia pallida* NF-κB (AIPGENE8848) from the ReefGenomics database, and *Orbicella faveolata* NF-κB (18), and *N. vectensis* (Nv) NF-κB, *Drosophila* Dif and Dorsal, and *Homo sapiens* p100, p105, RelA, RelB, and c-Rel from the UniProt database. The tree was rooted with RHD of *Capsaspora owczarzaki* NF-κB (from UniProt database). Conserved motifs from MEME analysis were truncated based on motif predictions (Supplemental Table 1), were aligned by Clustal Omega (35), and were inputted into PAUP* (36) for a distance neighbor-joining analysis bootstrapped 1000 times.

### Plasmid constructions, cell culture and transfections

Expression plasmids for FLAG-tagged human IKKβ, FLAG-Aq-NF-κB, FLAG-Nv-NF-κB, FLAG-Hu-IKKβ, and the empty pcDNA-FLAG vector have been described previously (17–19). The cDNA and PCR-generated Aq-NF-κB truncation mutants (Aq-RHD and Aq-Cterm) were subcloned into pcDNA-FLAG or the yeast GAL4-fusion vector pGBT9. Details about primers and plasmid constructions are included in Supplemental Tables 2 and 3, respectively.

### Cell culture and transfections

DF-1 chicken fibroblasts and human HEK 293 or 293T cells were grown in Dulbecco’s modified Eagle’s Medium (DMEM) (Invitrogen) supplemented with 10% fetal bovine serum (Biologos), 50 units/ml penicillin, and 50 μg/ml streptomycin as described previously (19). Transfection of cells with expression plasmids was performed using polyethylenimine (PEI) (Polysciences, Inc.) essentially as described previously (19,37). Briefly, on the day of transfection, cells were incubated with plasmid DNA and PEI at a DNA:PEI ratio of 1:6. Media was changed ∼20 h post-transfection, and whole-cell lysates were prepared 24 h later in AT Lysis Buffer (20 mM HEPES, pH 7.9, 150 mM NaCl, 1 mM EDTA, 1 mM EGTA, 20% w/v glycerol, 1% w/v Triton X-100, 20 mM NaF, 1 mM Na_4_P_2_O_7_· 10H_2_O, 1 mM dithiothreitol, 1 mM phenylmethylsulfonyl fluoride, 1 μg/ml leupeptin, 1 μg/ml pepstatin A, 10 μg/ml aprotinin). DF-1 cells analyzed by immunofluorescence were passaged onto glass coverslips on the day prior to fixation.

### Lysis of sponge tissue

Black encrusting sponge tissue was obtained from the Boston University Marine Program lab by scraping tissue off a rock substrate, immediately flash freezing the tissue, and storing at −80 °C until use. To prepare a sponge lysate, a piece of tissue approximately 2 cm by 2 cm was placed into a glass dounce tissue grinder (Wheaton USA) with 1 ml of AT Lysis Buffer and protease inhibitors (described above). Tissue was ground approximately ten times by hand, and then the sample was transferred to a 1.5-ml microcentrifuge tube and gently rotated for 1 h at 4 °C. The sample in buffer was then stored at −80 °C until use.

### Western blotting, electrophoretic mobility shift assays (EMSAs), reporter gene assays, and indirect immunofluorescence

Western blotting was performed essentially as described previously (17,19). Briefly, cell and tissue extracts were separated on a 10% SDS-polyacrylamide gels. Proteins were then transferred to nitrocellulose at 4°C at 250 mA for 2 h followed by 160 mA overnight. The membrane was blocked in TBST (10 mM Tris-HCl [pH 7.4], 150 mM NaCl, 0.1% v/v Tween 20) containing 5% powered milk (Carnation) for 1 h at room temperature. Filters were incubated at 4 °C with primary antiserum diluted in 5% milk TBST as follows: FLAG antiserum (1:1000, Cell Signaling Technology), α-tubulin antiserum (1: 1000, Cell Signaling Technology) or Ap-NF-κB antiserum (1:1000, (17)) After extensive washing in TBST, filters were incubated with anti-rabbit horseradish peroxidase-linked secondary antibody (1: 4000, Cell Signaling Technology) or anti-mouse horseradish peroxidase-linked secondary antibody (1:3000, Cell Signaling Technology). Immunoreactive proteins were detected with SuperSignal West Dura Extended Duration Substrate (Pierce).

EMSAs were performed using a ^32^P-labeled κB-site probe (GGGAATTCCC, see Supplemental Table 2) and 293T whole-cell extracts or sponge tissue lysates (see above), essentially as described previously (17,19). Yeast GAL4-site *LacZ* and 293 cell κB-site luciferase reporter gene assays were performed as described previously (19). Transfection and indirect immunofluorescence of DF-1 cells were performed essentially as described previously, with fixed cells that were probed with rabbit anti-FLAG primary antiserum (1:50, Cell Signaling Technology) (19).

### Indirect immunofluorescence of sponge tissue

Flash-frozen black encrusting sponge tissue was placed into ice-cold 4% paraformaldehyde in full-strength filtered artificial sea water (ASW) (Instant Ocean) to fix overnight at 4 °C. Fixed tissue was then washed three times with filtered ASW, and then dehydrated in 30% sucrose in ASW at 4 °C overnight. Samples were then embedded in Optimal Cutting Temperature Compound (Sakura Tissue-Tek), cryosliced on a microtome into 45 uM slices, placed onto Superfrost Plus Microscope Slides (Fisher Scientific), and kept at −20 °C until subjected to preparation for immunofluorescence. Tissue slices were rehydrated with room temperature PBS, and the tissue slices were then placed in wells of a 48-well plate using forceps. Tissue slices were then washed with agitation three times with 1X PBS, then blocked and permeablized with 0.3% Triton X-100 + 5% Goat Serum (Gibco) in PBS for 1 h at room temperature. Slices were then incubated with 0.3% Triton X-100 plus 5% Goat Serum and anti-Ap-NF-κB antibody (17) (1:5000) in PBS overnight at 4 °C in a humidified chamber, with rotation. Slides were then washed with agitation three times with 0.3% Triton X-100 in 1X PBS, and incubated with secondary antibody (goat-anti-rabbit Alexa Fluor 488 (1:500), and 5% Goat Serum/0.3% Triton X-100 in PBS. Slices were then washed three times with 0.3% Triton X-100 in PBS, and on the last wash Hoechst (1:5000 of 20 uM) was added for 10 min. Slices were then washed three more times with 0.3% Triton X-100 in PBS, and then mounted onto Superfrost Plus Microscope Slides (Fisher Scientific) with Prolong Gold (Molecular Probes by Life Technologies) with coverslips and imaged on a confocal microscope (Nikon C2 Si).

### In vitro *kinase assays*

*In vitro* kinase assays were performed as described previously (17–19). Briefly, human 293T cells were transfected with pcDNA-FLAG-IKKβ, and pcDNA-FLAG-Aq-TBK constructs, lysed two days later, and the kinases were immunoprecipitated with anti-FLAG beads (Sigma). The immunoprecipitates were then incubated with approximately 4 μg of GST alone, GST-Aq-NF-κB or GST-Aq-NF-κB-ALA C-terminal peptides and 5 μCi [γ-^32^P] ATP (Perkin Elmer) in kinase reaction buffer (25 mM Tris-HCl, pH 7.5, 20 mM β-glycerophosphate, 10 mM NaF, 10 mM MgCl_2_, 2 mM DTT, 500 μM Na_3_VO_4_, 50 μM ATP) for 30 min at 30°C. Samples were then boiled in 2X SDS sample buffer and electrophoresed on a 10% SDS-polyacrylamide gel. The ^32^P-labeled GST-Aq-NF-κB peptides were detected by phosphorimaging. As a control for protein input, 4 μg of GST alone, GST-Aq-NF-κB or GST-Aq-NF-κB-ALA was electrophoresed on a 10% SDS-polyacrylamide gel, and proteins were detected by staining with Coomassie blue (Bio-Rad).

### LPS stimulation of black encrusting sponge tissue

Black encrusting sponge tissue of approximately 1 cm × 1 cm was obtained from the Boston University Marine Program lab by collecting tissue off a rock substrate and transferring it immediately into a glass dish with ASW and allowing it to acclimate at 25°C with light provided by Sylvania Gro-Lux (GRO/Aq/RP) fluorescent bulbs at approximately 20 μmol photons/m_2_/sec for 1 h. The sample was then cut in half with a clean razor and each piece was placed in a 6-well plate with 10 ml of ASW and lipopolysaccharide (LPS) from *Escherichia coli* 0111:B4 (Sigma) at a final concentration of μg/ml (or an equal volume of sterile diH_2_O, which the LPS was dissolved in). The LPS or water was applied directly above the surface of the tissue and allowed to incubate for 30 min. The tissue samples were then immediately placed into AT buffer and lysed as described above before being subjected to Western blotting or EMSA.

## Author contributions

L.M.W. experimental design and execution; M.M.I. luciferase assays (Fig. 2D), IKK processing (Fig. 3C), and Fig. 6; K.M.M. EMSA (Fig. 2F); authors in BB522 design and construction pcDNA vectors and immunofluorescent analysis of proteins in DF-1 cells (Fig. 2B, Fig. 3A); L.M.W. and T.D.G. design and supervision of experiments and manuscript preparation.

## Acknowledgments

We thank Bernard Degnan for the Aq-NF-κB cDNA, Justin Scace for black encrusting sponge tissue, Todd Blute and Zachary Gardner for help with microscopy, and Joshua Aguirre, Milad Babaei, Joseph Brennan, Chris DiRusso, and Yuekun Liu for helpful discussions.

## FOOTNOTES

1 To whom correspondence should be addressed: Dr. Thomas D. Gilmore, Department of Biology, Boston University, 24 Cummington Mall, Boston, MA 02215, Tel: 617-353-3445; Fax: 617-353-6340;

2 The abbreviations used are: aa, amino acid(s); Aq, *Amphimedon queenslandica*; Ap, Aiptasia; ASW, artificial seawater; IKK, IkappaB kinase; LPS, lipopolysaccharide; NF-κB, nuclear factor kappa B; Nv, *Nematostella vectensis*.

