## Supplemental for "Transcription factor NF-κB in a basal metazoan, the sponge, has conserved and unique sequences, activities, and regulation"

*Running title: *Sponge NF-κB*

*To whom correspondence should be addressed: Thomas D. Gilmore, Department of Biology, Boston University, 24 Cummington Mall, Boston, MA 02215; Tel: 617-353-5444; Fax: 617-353-6340;

**Keywords:** NF-κB transcription factor, evolution, lipopolysaccharide, signal transduction, immunology, sponge, *Amphimedon queenslandica*

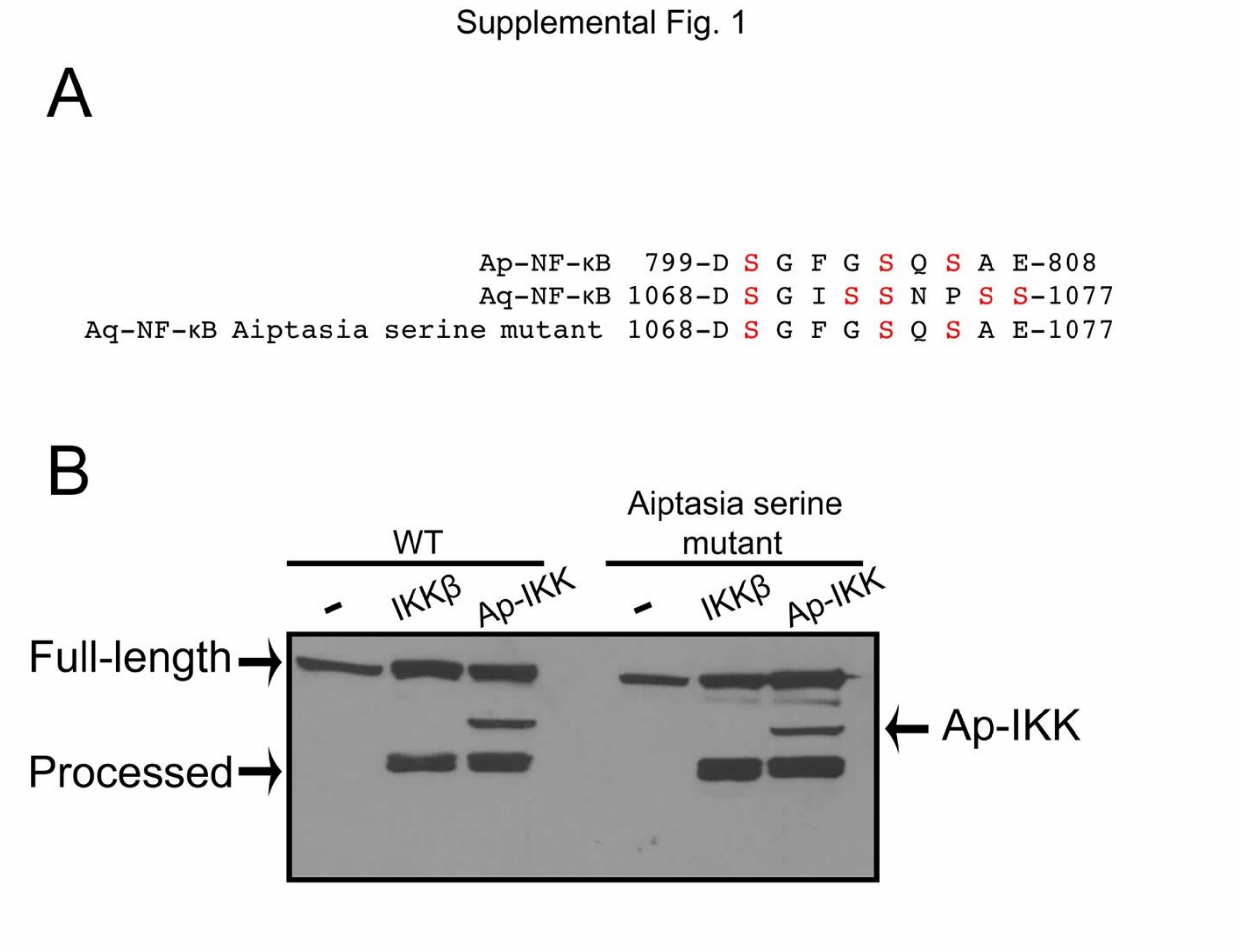


**Supplemental Figure 1. IKKs can induce processing in both WT Aq-NF-κB and Aq-NF-κB-Aiptasia serine mutant.** *A*, Amino acid (aa) sequence for processing and phosphorylation by IKKs in Ap-NF-κB (top) aligned with Aq-NF-κB aa sequence (middle). Aq-NF-κB-Aiptasia serine mutant (bottom) was created by changing Aq-NF-κB aa 1068-1077 to aa 799-808 from Ap-NF-κB. Shown in red are relevant serines. *B*, Western blot of lysates from 293T cells co-transfected with Aq-NF-κB wild-type (WT) or Aq-NF-κB-Aiptasia serine mutant and with empty vector (-) or HA-Hu-IKKβ or FLAG-Ap-IKK (bottom). Full-length and processed forms of Aq-NF-κB, and FLAG-Ap-IKK are indicated.

Supplemental Fig. 2

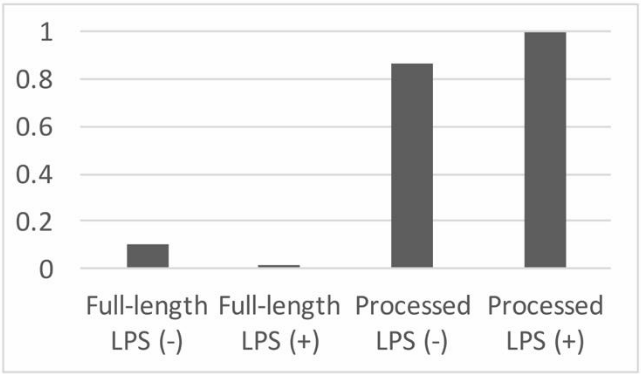


**Supplemental Figure 2. NF-κB processing is induced upon LPS stimulation in black encrusting sponge tissue.** The amount of NF-κB protein as compared to α-tubulin protein, as a percentage of processed NF-κB when stimulated with LPS.

**Supplemental Table 1. RHD sequences (bold) used for phylogenetic tree analysis.**

| Taxa: protein | Full-length NF-κB with Rel Homology Domain (RHD) in bold |
|---|---|
| ***Homo sapiens*: p100** | MESCYNPGLDGIIEYDDFKLNSSIVEPKEPAP<br>ETADG**PYLVIVEQPKQRGFRFRYGCEGPS**<br>**HGGLPGASSEKGRKTYPTVKICNYEGPAK**<br>**IEVDLVTHSDPPRAHAHSLVGKQCSELGI**<br>**CAVSVGPKDMTAQFNNLGVLHVTKKNM**<br>**MGTMIQKLQRQRLRSRPQGLTEAEQREL**<br>**EQEAKELKKVMDLSIVRLRFSAFLRASDG**<br>**SFSLPLKPVISQPIHDSKSPGASNLKISRMD**<br>**KTAGSVRGGDEVYLLCDKVQKDDIEVRF**<br>**YEDDENGWQAFGDFSPTDVHKQYAIVFR**<br>**TPPYHKMKIERPVTVFLQLKRKRGGDVS**<br>**DSKQFTYYPLVEDKEEVQRKRRKALPTFS**<br>**QPF**GGGSHMGGGSGGAAGGYGGAGGGGS<br>LGFFPSSLAYSPYQSGAGPMGCYPGGGGGA<br>QMAATVPSRDSGEEAAEPSAPSRTPQCEPQA<br>PEMLQRAREYNARLFGLAQRSARALLDYG<br>VTADARALLAGQRHLLTAQDENGDTPLHL<br>AIIHGQTSVIEQIVYVIHHAQDLGVVNLTNH<br>LHQTPLHLAVITGQTSVVSFLLRVGADPALL<br>DRHGDSAMHLALRAGAGAPELLRALLQSG<br>APAVPQLLHMPDFEGLYPVHLAVRARSPEC<br>LDLLVDSGAEVEATERQGGRTALHLATEME<br>ELGLVTHLVTKLRANVNARTFAGNTPLHLA<br>AGLGYPTLTRLLLKAGADIHAENEEPLCPLP<br>SPPTSDSDSDSEGPEKDTRSSFRGHTPLDLTC<br>STKVKTLLLNAAQNTMEPPLTPPSPAGPGLS<br>LGDTALQNLEQLLDGPEAQGSWAELAERLG<br>LRSLVDTYRQTTSPSGSLLRSYELAGGDLAG<br>LLEALSDMGLEEGVRLLRGPETRDKLPSTAE<br>VKEDSAYGSQSVEQEAEKLGPPPEPPGGLCH<br>GHPQPQVH |
| ***Homo sapiens*: p105** | MAEDDPYLGRPEQMFHLDPSLTHTIFNPEVF<br>QPQMALPTDG**PYLQILEQPKQRGFRFRYV**<br>**CEGPSHGGLPGASSEKNKKSYPQVKICNY**<br>**VGPAKVIVQLVTNGKNIHLHAHSLVGKH**<br>**CEDGICTVTAGPKDMVVGFANLGILHVT**<br>**KKKVFETLEARMTEACIRGYNPGLLVHP**<br>**DLAYLQAEGGGDRQLGDREKELIRQAAL**<br>**QQTKEMDLSVVRLMFTAFLPDSTGSFTRR**<br>**LEPVVSDAIYDSKAPNASNLKIVRMDRTA**<br>**GCVTGGEEIYLLCDKVQKDDIQIRFYEEE**<br>**ENGGVWEGFGDFSPTDVHRQFAIVFKTP**<br>**KYKDINITKPASVFVQLRRKSDLETSEPKP**<br>**FLYYPEIKDKEEVQRKRQKLMPNFSDSF**G<br>GGSGAGAGGGGMFGSGGGGGGTGSTGPGY |

| | |
|---|---|
| | SFPHYGFPTYGGITFHPGTTKSNAGMKHGT<br>MDTESKKDPEGCDKSDDKNTVNLFGKVIET<br>TEQDQEPSEATVGNGEVTLTYATGTKEESA<br>GVQDNLFLEKAMQLAKRHANALFDYAVTG<br>DVKMLLAVQRHLTAVQDENGDSVLHLAIIH<br>LHSQLVRDLLEVTSGLISDDIINMRNDLYQT<br>PLHLAVITKQEDVVEDLLRAGADLSLLDRL<br>GNSVLHLAAKEGHDKVLSILLKHKKAALLL<br>DHPNGDGLNAIHLAMMSNSLPCLLLLVAAG<br>ADVNAQEQKSGRTALHLAVEHDNISLAGCL<br>LLEGDAHVDSTTYDGTTPLHIAAGRGSTRL<br>AALLKAAGADPLVENFEPLYDLDDSWENA<br>GEDEGVVPGTTPLDMATSWQVFDILNGKPY<br>EPEFTSDDLLAQGDMKQLAEDVKLQLYKLL<br>EIPDPDKNWATLAQKLGLGILNNAFRLSPAP<br>SKTLMDNYEVSGGTVRELVEALRQMGYTE<br>AIEVIQAASSPVKTTSQAHSLPLSPASTRQQI<br>DELRDSDSVCDSGVETSFRKLSFTESLTSGA<br>SLLTLNKMPHDYGQEGPLEGKI |
| ***Homo sapiens*: RelA** | MDELFPLIFPAEPAQASG**PYVEIIEQPKQRG**<br>**MRFRYKCEGRSAGSIPGERSTDTTKTHPT**<br>**IKINGYTGPGTVRISLVTKDPPHRPHPHEL**<br>**VGKDCRDGFYEAELCPDRCIHSFQNLGIQ**<br>**CVKKRDLEQAISQRIQTNNNPFQVPIEEQ**<br>**RGDYDLNAVRLCFQVTVRDPSGRPLRLPP**<br>**VLSHPIFDNRAPNTAELKICRVNRNSGSCL**<br>**GGDEIFLLCDKVQKEDIEVYFTGPGWEAR**<br>**GSFSQADVHRQVAIVFRTPPYADPSLQAP**<br>**VRVSMQLRRPSDRELSEPMEFQYLPDTDD**<br>**RHRIEEKRKRTYETFKSIM**KKSPFSGPTDP<br>RPPPRRIAVPSRSSASVPKPAPQPYPFTSSLST<br>INYDEFPTMVFPSGQISQASALAPAPPQVLPQ<br>APAPAPAPAMVSALAQAPAPVPVLAPGPPQ<br>AVAPPAPKPTQAGEGTLSEALLQLQFDDED<br>LGALLGNSTDPAVFTDLASVDNSEFQQLLN<br>QGIPVAPHTTEPMLMEYPEAITRLVTGAQRP<br>PDPAPAPLGAPGLPNGLLSGDEDFSSIADMD<br>FSALLSQISS |
| ***Homo sapiens*: RelB** | MLRSGPASGPSVPTGRAMPSRRVARPPAAP<br>ELGALGSPDLSSLSLAVSRSTDELEIIDEYIKE<br>NGFGLDGGQPGPGEGLPRLVSRGAASLSTV<br>TLGPVAPPATPPPWGCPLGRLVSPAPGPGPQ<br>**PHLVITEQPKQRGMRFRYECEGRSAGSIL**<br>**GESSTEASKTLPAIELRDCGGLREVEVTA**<br>**CLVWKDWPHRVHPHSLVGKDCTDGICR**<br>**VRLRPHVSPRHSFNNLGIQCVRKKEIEAAI**<br>**ERKIQLGIDPYNAGSLKNHQEVDMNVVRI**<br>**CFQASYRDQQGQMRRMDPVLSEPVYDK**<br>**KSTNTSELRICRINKESGPCTGGEELYLLC**<br>**DKVQKEDISVVFSRASWEGRADFSQADVH**<br>**RQIAIVFKTPPYEDLEIVEPVTVNVFLQRL** |

| | |
|---|---|
| | **TDGVCSEPLPFTYLPRDHDSYGVDKKRKR**<br>**GMPDVLGEL**NSSDPHGIESKRRKKKPAILD<br>HFLPNHGSGPFLPPSALLPDPDFFSGTVSLPG<br>LEPPGGPDLLDDGFAYDPTAPTLFTMLDLLP<br>PAPPHASAVVCSGGAGAVVGETPGPEPLTL<br>DSYQAPGPGDGGTASLVGSNMFPNHYREA<br>AFGGGLLSPGPEAT |
| ***Homo sapiens*: c-Rel** | MASGAYN**PYIEIIEQPRQRGMRFRYKCEG**<br>**RSAGSIPGEHSTDNNRTYPSIQIMNYYGKG**<br>**KVRITLVTKNDPYKPHPHDLVGKDCRDG**<br>**YYEAEFGQERRPLFFQNLGIRCVKKKEV**<br>**KEAIITRIKAGINPFNVPEKQLNDIEDCDL**<br>**NVVRLCFQVFLPDEHGNLTTALPPVVSNP**<br>**IYDNRAPNTAELRICRVNKNCGSVRGGDE**<br>**IFLLCDKVQKDDIEVRFVLNDWEAKGIFS**<br>**QADVHRQVAIVFKTPPYCKAITEPVTVKM**<br>**QLRRPSDQEVSESMDFRYLPDEKDTYGN**<br>**KAKKQKTTLLFQKL**CQDHVETGFRHVDQ<br>DGLELLTSGDPPTLASQSAGITVNFPERPRPG<br>LLGSIGEGRYFKKEPNLFSHDAVVREMPTG<br>VSSQAESYYPSPGPISSGLSHHASMAPLPSSS<br>WSSVAHPTPRSGNTNPLSSFSTRTLPSNSQGI<br>PPFLRIPVGNDLNASNACIYNNADDIVGMEA<br>SSMPSADLYGISDPNMLSNCSVNMMTTSSD<br>SMGETDNPRLLSMNLENPSCNSVLDPRDLR<br>QLHQMSSSSMSAGANSNTTVFVSQSDAFEG<br>SDFSCADNSMINESGPSNSTNPNSHGFVQDS<br>QYSGIGSMQNEQLSDSFPYEFFQV |
| ***Capsaspora owczarzaki*: NF-κB** | MDLSELSGWDPNLSLQEHTANLLAMDDST<br>MAAMILHSDGLSLFRDMGLYNSVSTSLDISG<br>IPKFPPPPQQPQLAQAPPRRGHNQSSSSDSHS<br>TPSPGSVLFSPSPASQDMSLQSPELGLVGLTG<br>NSRLASTSEADDILLANILGHPVRSVSANTS<br>MVGLPDDLGFSPTTGMDLTITATSPATADSS<br>ASATAFPAASPIASPSVSTSSGPVTVGGTSAA<br>LAALTPDRINRLLSAAEAAAEGAATLVE**DL**<br>**LMVTEEPAQFARFRYMSEQRERSLAGEN**<br>**SFPTLMVNPKYARVVPEMALVTAVLVTK**<br>**MPDPHTGRQQKHWHHLGGIPAAPLEGP**<br>**QRIARFDNIAVIMDKANNKDKDKSKAPVR**<br>**SKDDQRCVRIMFELVFVSGNTQFYGRAIS**<br>**QPIYNAK**LAITKISHSSGPVTGGNEVIMLCS<br>KIRKGVTGVRMTDPTQWSVQAPSGSAWEL<br>NPQTLKADCNVPGANLFFHHQYAVVLTLPP<br>YHTQTITAPVTVRISILDTDDETESQYVEYTY<br>LPAEAAVRNAELAARKRRRDDSMRDFMDR<br>FDGSDGGNGSGSGRGNNGGHDGSDANNNG<br>RGGGGGSSSSKGGDEPFNFNSLIPMHQHKL<br>HQLALSTVRAVQGFAASGDARYLLALHRQL<br>LAAPNENGDSPLHTAVAQGNLRSTMALLPL<br>LAAEDLQSVNDMGETVLHSAVIEKRAAIAR |

| | |
|---|---|
| | LLLVAGADLGQSNARNFNRNSLHYLARHG<br>DRATAMAVFGVFGSAQAPPANTNTPAQAP<br>AGETKPKPADLRLLARIQAQAIKALLACELE<br>TGATPAHLAIRGGHWHVFEACAKLAASAPI<br>PKAAGSLLSMVAEKSSGHSLLHSCVLANNE<br>QAVRLLINLGASGNARDFGKNTPLHLAARQ<br>GHIGIAALLVEAGATLSLNAVSQTPLDVLTS<br>EGSGLSRDQLRALVAVLRGELKYADMRGR<br>PTLRMPTHAELHSTAAALTSASPGAVSLADF<br>YAGKKASRSPAPLGASSSLLSSTGASAAGAS<br>APTIAAVHAASATPVERTSMNNDDDYVLLE<br>KDAPYPVEQQPHGKRNKHSHHRFTRSSHGS<br>QDKDELKKDKDDPKKEKEPKELSKFTLKEA<br>FVDGTNFWELTRKFAGKKKMASASTGEME<br>PLSPERPLSPTNAGSGAASPFNQAKEQVSPG<br>AVPPTGLEKLVNKLMDASEATLSSQPAEAV<br>TPEQKLAEKLEKLGLAPASTTSAPPPHPKVA<br>ALNAQSVEDARKTSTHALYSVD |
| ***Amphimedon queenslandica*: NF-κB** | MAFNGIDPSSLPEALIRDTMNTLPTHIVNSNP<br>DSLNSSPYSSLTQVR**LEIVEQPKSRGFRFRY**<br>**DCEGQSHGGLPGENSEKNRRQKTYPTVH**<br>**LKGYRGRARVMVSLVTDSDPAMPHAHSI**<br>**VGKNAIDGRCVVEIGPETDMYAQFTSLGI**<br>**LHVTKKKVPEVLTRRLLQQTTPRGQMVD**<br>**QMEVVDVDMTTAQLTSEEQDEIHQQAQT**<br>**LAKSMNLSVVRLCFQAFLPDENGRYTIPI**<br>**DPVFSNKVYDSKAPSAGTLKICRLDRTSGS**<br>**VKGGDDVFLLCDKVQKNDIEVVFYEDKQ**<br>**ETTGGMQLQPWMAKGRFGPNDVHHQYA**<br>**IVFQTPTFYNQAIEHPVQVWIALKRPSDH**<br>**ETSEPKPFLYLPQEFDEERIGQKRRKKITH**<br>**FNNFFEGP**GGGGGAGGGAGGAGGNFFSRD<br>FNYGSGGGYNSGFNFFGGSSGGGGGSGGGS<br>ANNAGTGGGTTFSGGNTSAANMPVSVDSL<br>YSTLPPSSNQHIFAATATNPHYPHMRQQQPH<br>GLSFSNGGGRGMGGTMYSHAMDQQFMSTA<br>SGHLVSRTGAGGVTVKREPPDYYMDVERD<br>NVQPPLPALSEEGGTGGGNIMRPKDQLPPSQ<br>RGPGDSGKLLDVSSNIEDVESGYVEAERMD<br>SGLPTSMAAEPSQESTSSETEAQQALQALKD<br>RQQMAFEVCDRMFNALLAWATTKDIRYLL<br>AAQRSLTAVQNQEGDTALHLAIIHNHQDVV<br>LQLLDVLPQLPPTETPVVDCLNNFKQSPVHL<br>AVITRQHKVVQYLLKANANPLVSDRNGDTP<br>LHLACKYGFLQGIVPLLNRSTRINTEGCRIPE<br>LVMRNNDGLTPLHLAAACGNPDCFKELVK<br>AHADVNVQDSKSGKSALHYLIEKGDLPLTG<br>FLITESETNIECTDFSGNTPLHCAAALGNVAI<br>VSLLIAAGANLVCQNQEGELPLVLAEYGGH<br>EEVVKVLKDSLVKAGLDKPEEQLSTQMKSV<br>SLTEEDKALAALRANSSEGDLSKLDFRPRIS |

| | |
|---|---|
| | LALILDPINEGCDWKALAKCLSLSHLEAGLE<br>AMTSPTKELLTMYEACDGTIAKLRQALLDI<br>NRSDAVNIIDRYMQEEKGIVTSKQTYDSGIS<br>SNPSSLSNEQVVGKHSTIPSSQV |
| ***Nematostella vectensis:*** **NF-κB** | MAQSEQQVGSALTESMLNEIIRPGYLPDISA<br>LHVPLGTNAEEPSYTE**PYLEILEQPKPRGF**<br>**RFRYPSEGPSHGGLPGQFSTSKSKSYPSVQ**<br>**VNNYQGPCRIVVTLVTKDEPYMLHAHSL**<br>**TGKNANEEGVVTVQVGPDQHMTASFPNL**<br>**GIQHVTKKNVVKVLMDRFIKWQTLQNAT**<br>**FAKLSEGIKDGVDLSLFGVNTAINSNKLGF**<br>**DKNVALSVANQEAAKSREYAKQQAAAM**<br>**DLSAVRLCFQAYLPDQDGNFTRPLKPVYS**<br>**DAVLDSKEPSASQLKICRMDKNSGCVTGG**<br>**DEIYLLCDKVQKDDIEIHFYEMDDITGKY**<br>**TWEDLGKFSPCDVHRQFAIVFKTPPYWNI**<br>**AIERPANVLVELRRKKNGGETSEPVQFTY**<br>**QPQLFDKEAIGAKRRKTVPHFTEFLSGGS**<br>**SGATGGGGSSVSGFNFSADFL**QQGVFLTQ<br>NPSNM |
| ***Aiptasia pallida*****: NF-κB** | MTHSEQQVGGTLTDSMLHEIMTPGFLPDISS<br>LNVQMEMGYEG**PYLEILEQPKSRGFRFRY**<br>**PCEGPSHGGLPGEFSDSKNKSYPSVQVCN**<br>**YQGPCRIVVSLVTEDEPHMPHAHSLTGK**<br>**HANNDGIVTVQIGTEQGMTASFPNLGIQH**<br>**VTKKKVAKTLTERYTKMQALQNATLTAL**<br>**ATNNSTPSSFMNFGSVAREQVMASQGPFD**<br>**RNLAAAVAGEETKKILKLVQEQSKTMNL**<br>**SAVRLCFQAYLPDENGNFTKPLKPCISNP**<br>**VYDSKAPASCQLKICRMDKNSGCVTGGD**<br>**EIYLLCDRVQKDDIEIRFYENNDDGKPIW**<br>**EDTGKFAPADVHRQFAIVFKTPAYHNIAIE**<br>**RPVEVLLELRRKSDKETSEPFTFTYSPQM**<br>**FDTEQIGAKRQKKVPHFSDYYPPG**GPPGA<br>AGGGGGGFNFGSGFLFDFGFGGIPGVGASTS<br>TSTQQSQGSSSGTTQNAMSSTQPSQEMLKEL<br>AWAIAKHTSSAMQDYAATGDVRYLLSVQR<br>QITAVQNDDGDTALHIAIINCQFTAIEGLVSV<br>MKDLQGDFINTFNYLRQTPLQLATITKQALA<br>TECLLRGNADATLRDRHGNTPVHTACAQG<br>DVHCLRVLLDTKLRKEKDGFPELHWQNYD<br>GYTPLHLAVIKGNREIIQILLSEGANVESKDG<br>TCGRSPLHLAIEHDNLAISGYLILEARCDVDS<br>LTYDDNTPLHLAAGLGLVGETALLVAAGA<br>DTMATNSEDETPYSLATTAEVKKILGDDEG<br>PSDSITTSNTDPVITDIAEKVNKNVQRFTTPP<br>ESESLVNSSNYGDYKHWQNRQEMDREDSG<br>FGSQSAERDNDSLLLSLTTHPDKIMRPITDQE<br>PRFDIHHGQPMGRTN |
| ***Orbicella faveolata*****: NF-κB** | MATNSERQIGETLTDSLLMDIMTPGYLPDIS<br>ALQVPTATYSG**PYMEILEQPKQRGFRFRY** |

| | |
|---|---|
| | **PCEGPSHGGLPGQYSEKGKKSYPSVQLCN**<br>**YHGPARIVVSLVTVDEPPMPHAHSLIGKN**<br>**SNNGAVTVQIGPEHGMTASFPNLGIQHVT**<br>**KKSVGKVLMERYIKMQTLHTATLHALTA**<br>**DSKGFDMELVGDQALADGETATFNRTMA**<br>**EAVAAEESQKVRQMVEDQKQSMNLNAV**<br>**RLCFQAYLPDDGGCFTKALPPCISNPVYD**<br>**SKAPSASNLKICRMDRNSGCVTGGDEVYL**<br>**LCDKVQKEDIDVMFYEIDVETGKKTWEA**<br>**GGVFAPTDVHRQVAIVFKTPAYWNIATER**<br>**PVKVHLELRRKSDQETSEPVEFTYQPQLF**<br>**DKEQIGAKRRKKIPHFSDYF**SGGGGGGGP<br>PGMGGAGAGGGGFGTFAPLGLLGSALWFA<br>NPGTSQSNNQGGSGGAGTSQQGQSHSSGQS<br>QASGQSHQADLSELAWNLAEKSAAAMRDY<br>AATGDMRYLLAAQRHLTAVQDDSGDTALH<br>LAVINSQQEVVQCLIDIMAGLPESYISEYNFL<br>RQSPLHLAAITKQPRMLDCLLRASANVRSR<br>DRHGNTAVHIACMHGDAVCLKALLNYNVS<br>KTVLNWQNYQGVTPVHLAVLAGSKDVLKL<br>LNSAGANMSAQDGTSGKTPLHHAVEQDNL<br>AVAGFLILEANCDVDATTFDGNTPLHIAAAS<br>GLKGQTALLVAAGADTTLQNSDEETAFDLA<br>NVAEVQEILDEDEALSTDPSQDDELTAGLTG<br>LTLGQGDMDKLDPYVRRKMAQRLDPSIGA<br>DWRELAKRLGLGTLESAFAIHSSPTTQVLAQ<br>YEAADGSIKTLRQVLRDMRRGDVLEILDGR<br>KSPLSHDSGFDSGLGSQSLSAYRSEELASYE<br>AGSSSVPSSSSLQKGTTSLESTRKAGGYRPIR<br>QHQDVF |
| ***Drosophila*: Dif** | **MFEEAFGDIQEIINASMELNGGATGGGSV**<br>**AGAVGGGGAAHHILSQSTSLPVMPSHIPL**<br>**HLQNQNMNQNLPEPSARSGPHLRIVEEPT**<br>**SNIIRFRYKCEGRTAGSIPGMNSSSETGKT**<br>**FPTIEVCNYDGPVIIVVSCVTSDEPFRQHP**<br>**HWLVSKEEADACKSGIYQKKLPPEERRL**<br>**VLQKVGIQCAKKLEMRDSLVERERRNIDP**<br>**FNAKFDHKDQIDKINRYELRLCYQAFITV**<br>**GNSKVPLDPIVSSPIYGKSSELTITRLCSCA**<br>**ATANGGDEIIMLCEKIAKDDIEVRFYETD**<br>**KDGRETWFANAEFQPTDVFKQMAIAFKT**<br>**PRYRNTEITQSVNVELKLVRPSDGATSAPL**<br>**PFEYYPNPELLTKHNRRVAQK**TVESLKRS<br>LMSTNLHPSKQVKTSSQYTIFSKPQIATTTPQ<br>TQVSPGMPLMFPGGSPNFVQDIKMENGFMD<br>VDSQSSQCPSVERNFASPRSNCSTVDSIPPM<br>QMGQNQTHLYLPDATNFTFNGNFASPSSNC<br>STVDSIPPFQIGQRNNHMYLPENSNFPVNGC<br>SPTHFSGGSMTPINNNNNNVLINNNNNDFLSQ<br>KMSAISIPPQGNFGIKQVYQQTQQFLPQLQP |

| | |
|---|---|
| | ESIPYLAQSHPEQSQYQQQQQPQEQQPPADE<br>PTQSFSDLISSSIGMAPIDTSELIQDIEAELNS<br>LGIQPFK |
| ***Drosophila*: Dorsal** | MFPNQNNGAAPGQGPAVDGQQSLNYNGLP<br>AQQQQQLAQSTKNVRKK**PYVKITEQPAGK**<br>**ALRFRYECEGRSAGSIPGVNSTPENKTYPT**<br>**IEIVGYKGRAVVVVSCVTKDTPYRPHPHN**<br>**LVGKEGCKKGVCTLEINSETMRAVFSNL**<br>**GIQCVKKKDIEAALKAREEIRVDPFKTGF**<br>**SHRFQPSSIDLNSVRLCFQVFMESEQKGR**<br>**FTSPLPPVVSEPIFDKKAMSDLVICRLCSC**<br>**SATVFGNTQIILLCEKVAKEDISVRFFEEK**<br>**NGQSVWEAFGDFQHTDVHKQTAITFKTP**<br>**RYHTLDITEPAKVFIQLRRPSDGVTSEALP**<br>**FEYVPMDSGKHTFWNLHRHLK**RKPDEDL<br>FQQILRLDAKREVQPPTIEVIDLDTPKIDVQR<br>EIPSEMEFNHEESQQSEPALEQEQSVQQEQY<br>TQEQSLQQEQYTQEQSLQQEQYLQQLEQQQ<br>SFQLEEPMQQDQELPAQQSFDQAIDHLPDHT<br>SDHIPEDMEAADAHAEAEAHRLRSEQEKEI<br>DTIIDEKVRELEQLDLGQQLEPRPLTANDKIT<br>EWMKSSEIEQQVHEPSPTAEADVLDSALEIS<br>KADKTLDELLETVAELDEIYTDFKVQRDTY<br>KNTIQNELAGLQGRAPLQVEDSFDDAATYT<br>SLQIAFKNPVLIPMDDIMPPTPPMSQCAPED<br>AHQHYDPVEVNSQARKPETPMRPVPPVPPAI<br>LTIQYPPEEDKLPPLPPKRIRKQDSNAENRSI<br>EANTVQTKPSTGESPLNKRLPPAPKNPNFNT<br>LPRQKKPGFFSKLFSRRKSKPDLAQGQEN<br>SSILDSKANSREPSIGHFNMQDPMRASLRSS<br>KSAAPFISNPAPAKSSPVKAKKPGSKLTK<br>PVGRSVSSVSGKRPAYLNADVVHIPLKGDS<br>VNSLPQQQRTEGYSQSSTISVGAGLDRRTA<br>SALQLADIPISEGGMELVAIADRQSLHNLVS<br>SIEGHFNVQLDPNLDLTEAEHFALYTSIP<br>PLAAASEFDETSAYYAPVDAGEILTPDEVAK<br>RLAAANGI |

**Supplemental Table 2. List of primers used in these experiments.**

Primers for Subcloning (restriction enzyme sites used for subcloning are underlined)

| Primer Name | Primer Sequence |
|---|---|
| pcDNA-FLAG-Aq-RHD-F | 5’- CGAAGGATCCCTGCTTTTAATGGTATTGATCCC-3’ |
| pcDNA-FLAG-Aq-RHD-R | 5’-ATATCTCGAGCTAGTTCCCACCGGAGAAC -3' |
| pcDNA-FLAG-Aq-Cterm-F | 5’- ATCTAGAATTCAACTGGAGGCGGGACCACG -3’ |
| pcDNA-FLAG-Aq-Cterm-R | 5’- CGAGGGATCCCTAAACTTGACTAGAAGG -3' |
| GB-Aq-RHD-F | 5’-CGAAGGATCCCTGCTTTTAATGGTATTGATCCC-3’ |
| GB-Aq-RHD-R | 5’-ATATCTCGAGCTAGTTCCCACCGGAGAAC-3' |
| GB-Aq-Cterm-F | 5’-ATCTAGAATTCACTGGAGGCGGGACCACG-3’ |
| GB-Aq-Cterm-R | 5’-CGAGGGATCCCTAAACTTGACTAGAAGG-3' |
| Aq-NF-κB-ALA-F-1 | 5'-GCACTCCTGGATATCAATCGTA-3' |
| Aq-NF-κB-ALA-F-2 | 5'-CTTATGATGCTGGCATCGCTGCAAATCCTGCAGCTCTCTCTAATGAACAAG-3' |
| Aq-NF-κB-ALA-R-3 | 5'-CTTGTTCATTAGAGAGAGCTGCAGGATTTGCAGCGATGCCAGCATCATAAG-3' |
| Aq-NF-κB-ALA-R-4 | 5'-GCTCGAGCGGCCGCGCTCGAGGGATCCTAAACTTGACTAGAAGGAATGG-3' |
| Aq-NF-κB-DEL-F | 5'-CGCGGCTGCAGGACAGCAGTACAGAATC-3' |
| Aq-NF-κB-DEL-R | 5'-CGGGCCTGCAGTCCTCCCATTCCCCTC-3' |
| GST-Aq-NF-κB-Cterm-F | 5'-CGCGCGAATTCTAAATATAATTGATCGATATATGCAAG-3' |
| GST-Aq-NF-κB-Cterm-R | 5'-GCGCGCTCGAGCTACTTCCCTACAACTTGTTC-3' |
| Aq-NF-κB-SER-F-2 | 5'-GGCTTCGGTTCACAATCTGCAGAGCTCTCTAATGAACAAGTTGTAGG-3' |
| Aq-NF-κB-SER-R-3 | 5'-GAGCTCTGCAGATTGTGAACCGAAGCCAGAATCATAAGTCTGTTTAC-3' |

Primers for EMSA

| | |
|---|---|
| NF-κB-Consensus | 5’- TCGAGAGGTCGGGGAATTCCCCCCCCG -3’ |
| | 5’- TCGACGGGGGGGAATTCCCCGACCTC -3’ |

**Supplemental Table 3. List of plasmids used in these experiments.**

Expression Vectors for Use in Tissue Culture Cells and Yeast

| Plasmid Name | Plasmid Description |
|---|---|
| pcDNA-FLAG | pcDNA with a 5' FLAG Tag (19). |
| pcDNA-FLAG-Nv-NF-κB | Reference 19. |
| pcDNA-FLAG-Aq-NF-κB | BamHI-BamHI fragment containing amino acids 2-1096 of Aq-NF-κB was PCR-aimplified from pMD5-Aq-NF-κB and subcloned into Bam HI-digested pcDNA-FLAG (17). |
| pcDNA-FLAG-Aq-NF-κB-ALA | EcoRV-Aq-NF-κB-1037-1082-ALA and Aq-NF-κB-1066-1096-ALA-NotI PCR fragments were used as a template for assembly PCR of an EcoRV-NotI fragment containing full-length pcDNA-FLAG-Aq-NF-κB. Primers used for amplification were Aq-NF-κB-ALA-F-1, Aq-NF-κB-ALA-F-2, Aq-NF-κB-ALA-R-3, Aq-NF-κB-ALA-R-4. |
| pcDNA-FLAG-Aq-NF-κB-Δ | BamHI-PstI digested pcDNA-FLAG-Aq-NF-κB PCR product containing codons 2-511 and PsI-BamHI digested pcDNA-FLAG-Aq-NF-κB product containing codons 681-1096 were used as a template for assembly PCR of the two PCR products. Primers used for amplification were pcDNA-FLAG-Aq-RHD-F, pcDNA-FLAG-Aq-Cterm-R, Aq-NF-κB-DEL-F, Aq-NF-κB-DEL-R |
| pcDNA-FLAG-Aq-NF-κB-ΔALA | BamHI-PstI digested pcDNA-FLAG-Aq-NF-κB-ALA PCR product containing codons 2-511 and PsI-BamHI digested pcDNA-FLAG-Aq-NF-κB-ALA product containing codons 681-1096 were used as a template for assembly PCR of the two PCR products. Primers used for amplification were Aq-NF-κB-ALA-F-1, Aq-NF-κB-ALA-F-2, Aq-NF-κB-ALA-R-3, Aq-NF-κB-ALA-R-4. |
| pcDNA-FLAG-Aq-RHD | BamHI-XhoI digested Aq-RHD PCR product containing codons 2-452 was subcloned into BamHI- XhoI digested pcDNA-FLAG. Primers: pcDNA-FLAG-Aq-RHD-F and pcDNA-FLAG-Aq-RHD-R. PCR- amplified from pcDNA-FLAG-Aq-NF-κB. |
| pcDNA-FLAG-Aq-Cterm | EcoRI-BamHI digested Aq-Cterm PCR product containing codons 442-1095 was subcloned into EcoRI- BamHI digested pcDNA-FLAG. Primers: pcDNA-FLAG-Aq-Cterm-F and pcDNA-FLAG-Aq-Cterm-R. PCR- amplified from pcDNA-FLAG-Aq-NF-κB. |
| pcDNA-FLAG-Ap-IKK | Reference 17. |

| | |
|---|---|
| FLAG-Hu-IKKβ | Reference 38. |
| GBT9 | Reference 19. |
| GB-Nv-NF-κB | Reference 19. |
| GB-Aq-RHD | BamHI-XhoI digested Aq-RHD PCR product containing codons 2-452 was subcloned into BamHI- XhoI digested pcDNA-FLAG. Primers: GB-Aq-RHD-F and GB-Aq-RHD-R. PCR- amplified from pcDNA-FLAG-Aq-NF-κB. |
| GB-Aq-Cterm | EcoRI-BamHI digested Aq-Cterm PCR product containing codons 442-1095 was subcloned into EcoRI- BamHI digested pcDNA-FLAG. Primers: GB-Aq-Cterm-F and GB-Aq-Cterm-R. PCR- amplified from pcDNA-FLAG-Aq-NF-κB. |
| pcDNA-FLAG-Aq-NF-κB-Aiptasia serine mutant | EcoRV-Aq-NF-κB-1037-1082-SER and Aq-NF-κB-1066-1096-SER-NotI PCR fragments were used as a template for assembly PCR of an EcoRV-NotI fragment containing full-length pcDNA-FLAG-Ap-NF-κB. Primers used for amplification were Aq-NF-κB-ALA-F-1, Aq-NF-κB-SER-F-2, Aq-NF-κB-SER-R-3, Aq-NF-κB-ALA-R-4. |

Bacterial Expression Vectors

| **Plasmid Name** | **Plasmid Description** |
|---|---|
| pGEX-KG | Expression plasmid containing a 5' GST tag. |
| pGEX-KG-Aq-Cterm | EcoRI-XhoI fragment containing Aq-NF-κB codons 1048-1086 was subcloned into EcoRI-XhoI digested pGEX-KG. Primers GST-Aq-NF-κB-Cterm-F and GST-Aq-NF-κB-Cterm-R were used to PCR- amplify the fragment from pcDNA-FLAG-Aq-NF-κB. |
| pGEX-KG-Aq-Cterm-ALA | EcoRI-XhoI fragment containing Aq-NF-κB-ALA codons 1048-1086 was subcloned into EcoRI-XhoI digested pGEX-KG. Primers GST-Aq-NF-κB-Cterm-F and GST-Aq-NF-κB-Cterm-R were used to PCR- amplify the fragment from pcDNA-FLAG-Aq-NF-κB-ALA. |
